# Tissue-resident, extravascular CD64^-^Ly6C^-^ population forms a critical barrier for inflammation in the synovium

**DOI:** 10.1101/2021.02.25.432850

**Authors:** Anna B. Montgomery, Shang Yang Chen, Gaurav Gadhvi, Maximilian G. Mayr, Carla M Cuda, Salina Dominguez, Hadijat-Kubura Moradeke Makinde, Alexander Misharin, Miranda G. Gurra, Mahzad Akbarpour, Ankit Bharat, G. R. Scott Budinger, Deborah R. Winter, Harris Perlman

## Abstract

Monocytes are one of the most abundant immune cells infiltrating the inflamed organs. However, the majority of studies on monocytes focus on circulating cells, rather than those in the tissue. Here, we identify and characterize an intravascular (i.v.) and extravascular (e.v.) synovial population (Syn Ly6C^-^ cells) which lack cell surface markers of classical monocytes (Ly6C and CD62) or tissue macrophages (CD64 and Tim4), are transcriptionally distinct and conserved in RA patients. e.v. Syn Ly6C^-^ cells are independent of NR4A1 and CCR2, long-lived and embryonically derived while the i.v. Syn Ly6C^-^ cells are dependent on NR4A1, short lived and derived from circulating NCM. e.v. Syn Ly6C^-^ cells undergo increased proliferation and reverse diapedesis dependent on LFA1 in response to arthrogenic stimuli and are required for the development of RA-like disease. These findings uncover a new facet of mononuclear cell biology and are imperative to understanding tissue-resident myeloid cell function in RA.

## Introduction

In recent years, our understanding of the mononuclear phagocyte system has expanded, highlighting previously unknown complexities in cell origin and function. However, to date few studies have examined a role for monocytes in tissues, with the majority of studies centered on circulating monocytes, or monocyte-derived macrophages. Circulating monocytes exist in 3 main states, characterized by CCR2, CX3CR1, CD43 and Ly6c in mice: classical (CM) (CCR2^+^CX3CR1^low^CD43^-^Ly6c^hi^), intermediate (IM) (CCR2^±^CX3CR1^low^CD43^±^Ly6c^int^), and non-classical (NCM) (CCR2^-^CX3CR1^hi^CD43^+^Ly6c^low^) (Yona and Jung, 2010; Ziegler-Heitbrock et al., 2010). CM require CCR2 to exit the bone marrow, while NCM utilize sphingosine-1-phosphate receptor 5 (S1PR5) and/or CX3CR1 (Guilliams et al., 2018). Consequently, CCR2^-/-^ mice have reduced numbers of CM in circulation, while S1PR5^-/-^ and CX3CR1^-/-^ mice have reduced NCM (Guilliams et al., 2018). NCM also require CEBP/β for transcriptional activation of NR4A1 and CSF1R to maintain survival (Mildner et al., 2017). As such, NR4A1^-/-^ and CEBP/β mice also display markedly reduced numbers of circulating NCM. While transcriptional studies have exposed critical gene signatures for CM and NCM in the bone marrow and circulation, no such studies examined heterogeneity and function at the tissue level.

In contrast to well-characterized inflammatory CM (Guilliams et al., 2018), the direct impact of NCM in steady state and inflammation is unclear. The current dogma for circulating NCM centers on barrier maintenance due to the ability of NCM to adhere and patrol the endothelium (Narasimhan et al., 2019). In this context, NCM maintain the endothelium, scavenge debris, and elicit removal of damaged endothelial cells by neutrophils (Narasimhan et al., 2019). To date, only one study has proposed the existence of an NCM population in tissue (Schyns et al., 2019). The investigators identified a CD64^+^CD16.2^+^ subpopulation among extravascular CD45^+^Ly6c^lo^ cells in the lung that are derived from circulating NCM and require NR4A1. These cells were considered monocytes but were putative precursors for interstitial macrophages. It is not clear whether similar populations are found in other tissues due to the dearth of studies that have identified NCM in tissue that are distinct from circulating NCM.

It is established that recruitment of monocytes to the inflamed synovium is a requisite for sustainment and progression of rheumatoid arthritis (RA) (Udalova et al., 2016). Support for a functional role for NCM in RA comes from murine models. While complete ablation of circulating monocytes using clodronate-laden liposomes (Clo-lip) prevents the effector phase of K/BxN serum transfer-induced arthritis (STIA), pathology is restored exclusively with transfer of NCM, not CM (Misharin et al., 2014; Solomon et al., 2005). CX3CR1^-/-^ mice also display a reduction in STIA (Jacobs et al., 2010). In contrast, depletion of CM via anti-CCR2 antibody or deficiency in CCR2 has no effect on arthritis development in STIA (Misharin et al., 2014), TNF α -Transgenic mice (Puchner et al., 2018), or collagen-induced arthritis (Quinones et al., 2005). However, NR4A1^-/-^ mice remain sensitive to STIA and CIA regardless of reduced numbers of circulating NCM (Brunet et al., 2016; Liebmann et al., 2018). Taken together, these studies present a quandary on the role of NCM play RA.

To distinguish differential and distinct functional roles of monocytes, we focused on identifying the heterogeneity of CD64^-^Ly6C^-^ cells in synovial tissue. We uncovered two novel subpopulations of synovial Ly6C^-^ cells, which lack known markers of macrophages and can be separated by their intra-and extra-vascular location in mouse and human synovium. While intra-vascular synovial Ly6C^-^ cells retain a similar phenotype to PB NCM, the extra-vascular synovial Ly6C^-^ population are transcriptionally distinct from circulating NCM, independent of NR4A1 and CCR2, long-lived with embryonic origins, double in number within 24 hrs. post-arthrogenic insult and require reverse diapedesis mediated by LFA for the development of inflammatory arthritis. These data document an essential role for a newly described tissue-resident synovial Ly6C^-^ cell that functions as a physical barrier determining the susceptibility of the synovium to inflammatory arthritis.

## Results

### Non-classical monocytes in the synovium are distinct from those in the circulation

We sought to determine the contribution of circulating NCM to inflammatory arthritis by inducing STIA in NR4A1^-/-^ mice, which are depleted of NCM in peripheral blood (PB) (Figure 1A, Figure S1A-B). NR4A1^-/-^ mice developed STIA of comparable severity and onset to C57Bl/6 controls (Figure 1B), in agreement with a previous report (Brunet et al., 2016). Flow cytometry was then performed to identify monocyte populations preserved in the synovium of NR4A1^-/-^ mice, which may explain the sensitivity of these mice to inflammatory arthritis. We identified a novel synovial myeloid niche defined as CD45^+^CD11b^+^Ly6G^-^SigF^-^CD64^-^ of which the majority were Ly6C^-^. Based on this gating strategy synovial macrophages (CD64^+^) were the most abundant myeloid population in the synovium (Figure S1C), while Ly6C^-^ represented 10% of the synovial mononuclear phagocyte compartment, and Ly6C^+^ and Ly6C^int^ cells composed less than 1%. Surprisingly, synovial Ly6C^-^ (Syn Ly6C^-^) cell numbers remained unchanged in NR4A1^-/-^ mice (Figure 1C) even while NCM were markedly reduced in PB. In order to confirm that Syn Ly6C^-^ cells were not dependent on CCR2, STIA was induced in CCR2^-/-^ mice lacking CM in PB (Figure 1D, Figure S1B). CCR2^-/-^ mice also showed comparable clinical scores in STIA to C57Bl/6 controls (Figure 1E) as reported in previous studies (Jacobs et al., 2010; Misharin et al., 2014) and their numbers of Syn Ly6C^-^ cells were unchanged compared to controls (Figure 1F). Taken together, these data confirm that neither subtype of circulating monocyte is required for inflammatory arthritis, while a newly identified population of Syn Ly6C^-^ cells is independent of NR4A1 and CCR2, lacks expression of macrophage associated markers and may play an essential role in STIA.

**Figure 1:**
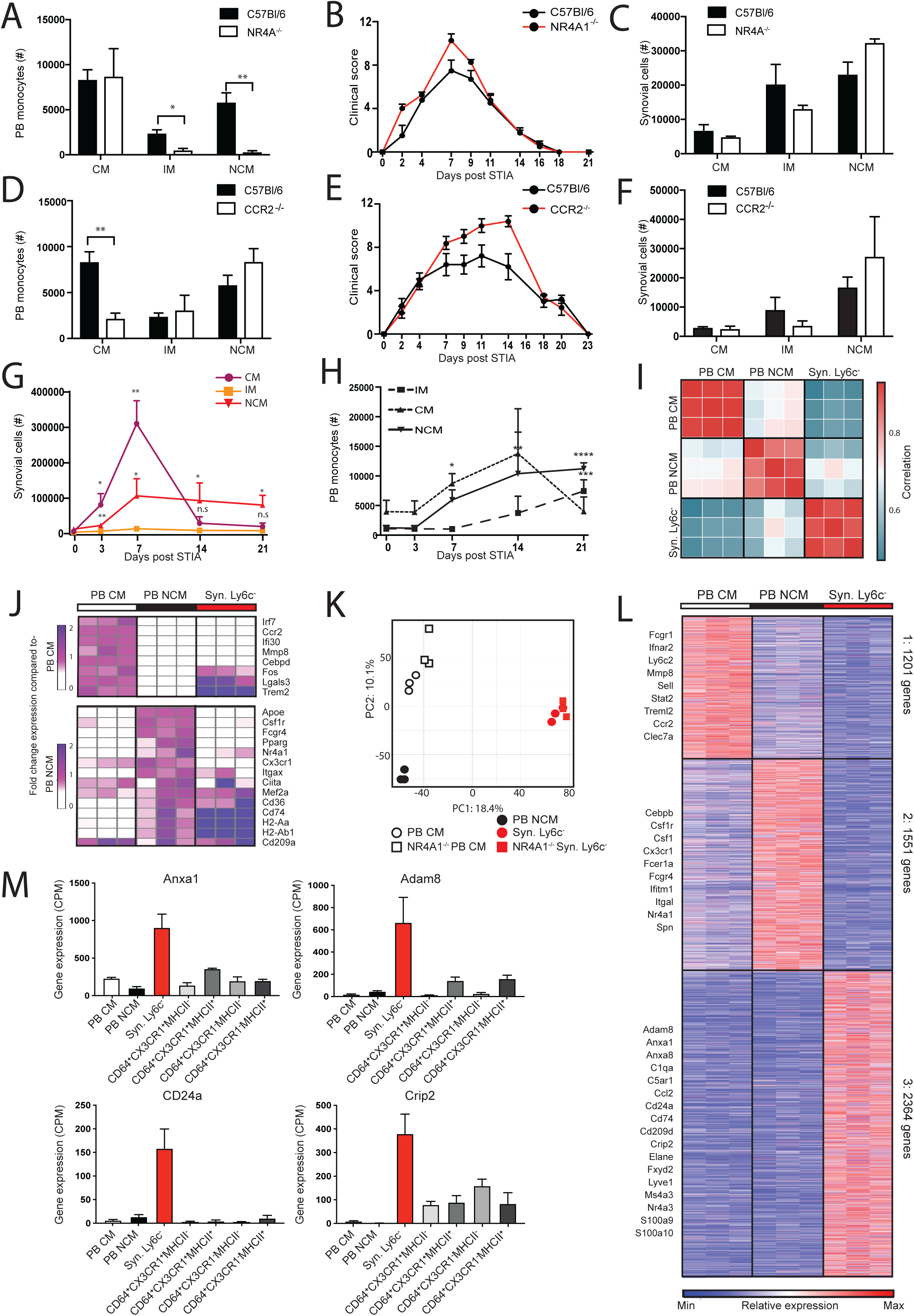
Synovial Ly6C^-^ are phenotypically distinct from circulating NCM. (A) Numbers of classical (CM), intermediate (IM), and non-classical (NCM) monocytes in the peripheral blood (PB), B) STIA severity and C) numbers of CM, IM, and NCM in synovium of C57Bl/6 compared to NR4A1^-/-^ mice, and in C57Bl/6 compared to CCR2^-/-^ mice (D-F). G) Changes in numbers of synovial Ly6C^+^, Ly6C^int^, and Ly6C^-^, and H) PB CM, IM, and NCM during STIA. Data shown are n>=4 ±SEM, *= P<0.05, **= P<0.01, ****=P<0.001. I) Pairwise Pearson’s correlation of global gene expression between replicates of PB CM, PB NCM and Syn Ly6C^-^. J) Fold-change expression of monocyte-associated genes from Mildner et al. compared to PB CM or NCM (Mildner et al., 2017). K) PCA of 10206 genes expressed by PB CM, PB NCM and Syn Ly6C^-^ from C57Bl/6 and NR4A1^-/-^ mice. L) k-means clustering of 5115 differentially expressed genes across PB CM, PB NCM and Syn Ly6C^-^. M) Mean expression of representative genes from PB CM, PB NCM, Syn Ly6C^-^ and Syn Mac populations (RNA-seq data: n=3, error bars indicate SEM).

Cell numbers were measured throughout disease to determine the response of synovial tissue myeloid populations during STIA. Synovial Ly6C^+^ and Ly6C^-^ numbers significantly expanded on D3 (p=0.003, p=0.006) and D7 (p=0.003, p=0.03) post-STIA compared to D0, while Ly6C^int^ cells were not significantly different at any timepoint (Figure 1G). By D14, synovial Ly6C^+^ cells returned to baseline whereas Ly6C^-^ cells plateaued on D14 and D21 compared to D0 (p= 0.05, p=0.02). Similarly, PB CM reach a peak prior to 21 days post-STIA while PB NCM continue to increase (Figure 1H).

To determine if Syn Ly6C^-^ cells exhibit a distinct transcriptional state from PB monocytes, we isolated CM and NCM from PB (PB CM, PB NCM) and Syn Ly6C^-^ cells from hindjoints of mice for RNA sequencing (RNA-seq). Given PB IM are likely an intermediate cell state, these cells were excluded from our studies. PB CM, PB NCM and Syn Ly6C^-^ cells exhibited distinct transcriptional profiles from each other (Figure 1I, Figure S1D). We then compared expression of genes associated with PB CM vs. PB NCM as described (Mildner et al., 2017). Expression of monocyte genes in PB CM and PB NCM largely aligned with expectations, but Syn Ly6C^-^ cells did not uniformly express PB CM genes – such as Irf7, Ccr2, Ifi30, Mmp8, and Cebpd – or those associated with PB NCM – such as Apoe, Csf1r, Fcgr4, Pparg, Nr4a1, and Cx3cr1 (Figure 1J). Furthermore, loss of NR4A1 had a minimal effect on the transcriptional profile of synovial Ly6C^-^ cells compared to WT (Figure1K, Figure S1E and supp. gene tables). These data show that Syn Ly6C^-^ cells are transcriptionally distinct from circulating cells and are not NR4A1 dependent.

K-means clustering of 5116 differentially expressed genes identified 3 gene clusters preferentially expressed by PB CM, PB NCM, or Syn Ly6C^-^ cells (Figure 1L, Supp. Table 1A). Compared to other clusters, the Syn Ly6C^-^ cell cluster (cluster 3) was enriched for genes associated with extracellular matrix organization, hormone secretion, cell division, cell adhesion, and regulation of biological processes (Figure S1F, Supp. Table 1B). Additionally, genes involved in antigen presentation (H2-Aa, Cd74), immune activation (Cd9, Pf4, Cd36), cell-to-cell interaction (Adam8, Anxa8) and cell differentiation (Cd24a, Crip2) were increased in Syn Ly6C^-^ cells compared to PB CM and PB NCM (Figure 1L). We found that on the individual gene and global level, Syn Ly6C^-^ cells exhibited a distinct transcriptional profile from each of the four CD64+ synovial macrophage populations (Syn Mac) (Figure 1M, Figure S1G-H, see Methods). Taken together, these data uncover a novel CD64-Syn Ly6C^-^ population present in the synovium.

### Single-cell RNA-sequencing identifies novel synovial Ly6C^-^ population

We utilized single-cell RNA-seq (scRNA-seq) to investigate heterogeneity of Syn Ly6C^-^ (CD45^+^CD11b^+^Ly6G^-^SigF^-^Tim4^-^CD64^-^). We defined 6 clusters (0-5) of myeloid subpopulations using unsupervised graph-based clustering of 7160 cells sequenced from C56BL/6 mice (Figures 2A, S2A,B, Supp. Table 2A). The expression of Cd14 and Itgam (CD11b) confirmed that the sorted cells were of myeloid lineage, while low expression of Fcgr1 (CD64), Timd4 (Tim4) and Mertk confirms non-macrophage classification (Figures 2B, S2C). Using singleR and reference population ImmGen dataset we found that subpopulations 0-3 exhibited a mixture of monocyte, macrophage, and DC annotations expected of tissue-resident myeloid cells, while subpopulations 4 and 5 were primarily assigned to monocytes and DC respectively (Figure 2C). Subpopulation 4 (tissue monocytes) also displayed similarity to PB CM and PB NCM (Figures 2D, S2B, S2D-F), while subpopulation 5 had the highest expression of genes associated with conventional DCs (Kirkling et al., 2018) (Figures 2E, S2G). Since subpopulation 2 expressed high levels of genes indicative of cell cycle (Figure S2H-J), our subsequent analysis centered on subpopulations 0, 1 and 3 which best reflected steady-state, tissue-resident, non-macrophage synovial cells.

**Figure 2:**
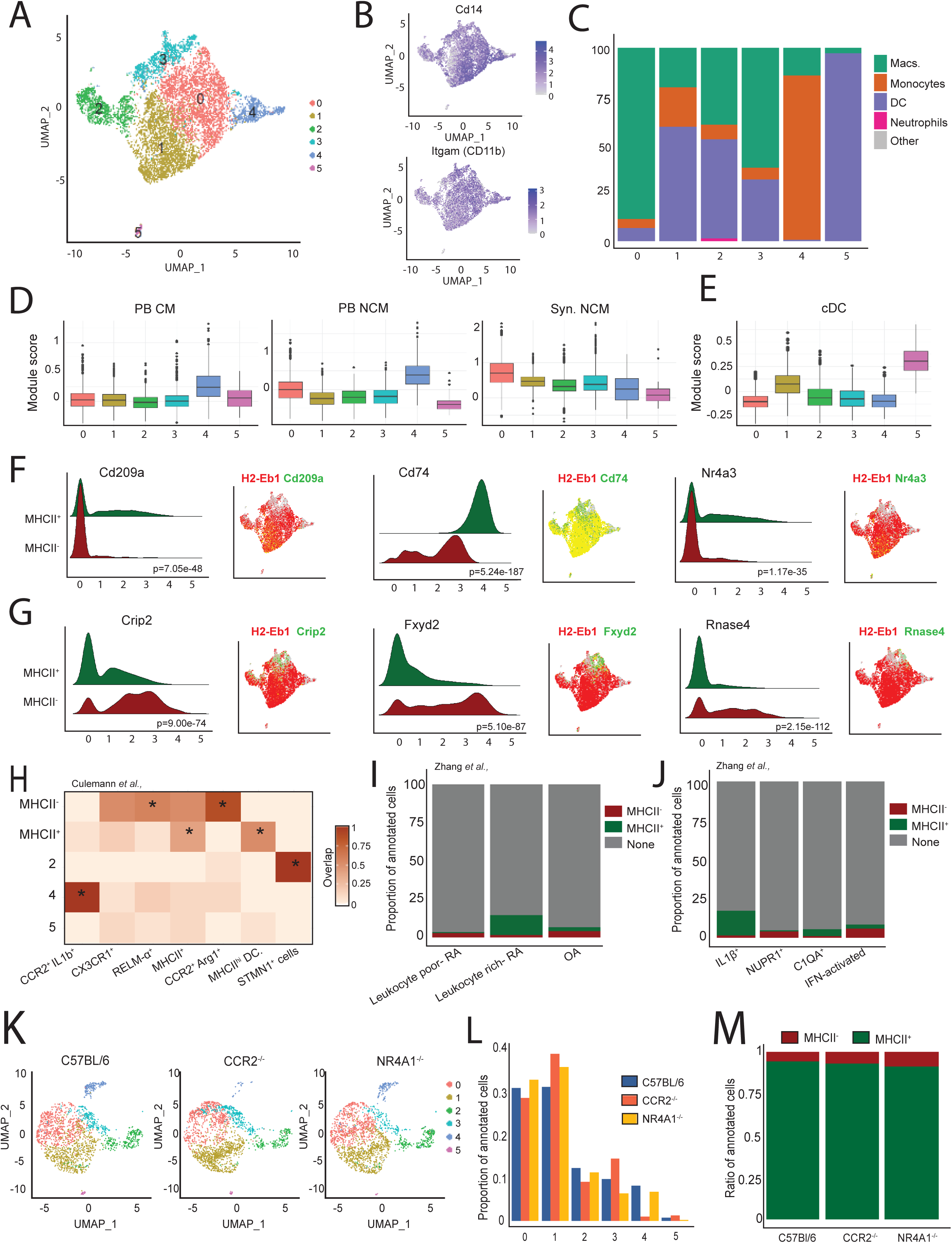
Single-cell RNA-sequencing analysis of joint myeloid niche identifies tissue Syn Ly6C^-^ cells. A) UMAP depicting 6 subpopulations of CD45^+^CD11b^+^Tim4^-^CD64^-^ cells from single-cell RNA-seq data. B) Expression of myeloid markers Cd14 and Itgam. C) Percent of cells in each subpopulation assigned to ImmGen cell types by singleR. D) Module score for each scRNA-seq subpopulation representing expression of key genes in PB CM, PB NCM, Syn. Ly6c^-^ and E) cDC. F-G) Ridge plots and UMAP visualization of gene expression by MHCII compartment. P-value by Wilcoxon test is indicated. H) Fraction overlap of differentially expressed genes from C57Bl/6 scRNA-seq subpopulations with top 20 markers of myeloid populations from murine synovium at day 5 post-STIA identified in (Culemann et al., 2019) * indicates significant p-value by hypergeometic test after FWER correctiron. I) Proportion of synovial myeloid cells annotated as MHCII+ or MHCII-over total cells from Leukocyte-poor RA, Leukocyte-rich RA, OA patients, J) and by cluster defined in published AMP data (Zhang et al., 2019). K) Integration of single-cell RNA-seq data on CD45^+^CD11b^+^Tim4^-^CD64^-^ cells from CCR2^-/-^ and NR4A1^-/-^ mice with C57Bl/6 (sub-sampled to 2000 cells). L) Proportion of cells annotated as each subpopulation in C57Bl/6, CCR2^-/-^ and NR4A1^-/-^ mice. M) Ratio of cells annotated as either MHCII^+^ or MHCII^-^ in C57Bl/6, CCR2^-/-^ and NR4A1^-/-^ mice.

Although subpopulations 0, 1 and 3 all had high similarity to Syn Ly6C^-^ cells from Figure 1 (Figures 2D, S2E), the variability in gene expression profiles suggested they may be distinct subtypes. In particular, subpopulation 1 exhibited higher expression of certain genes associated with the DC lineage (Figures 2E, S2B). Since MHCII genes (H2-Eb1, Ab1, Aa, DMb1, DMa) are associated with DCs, we partitioned subpopulations 0, 1 and 3 based on their expression, specifically on H2-Eb1 (Figure S2K-L). As expected, the MHCII^+^ compartment exhibited elevated expression of genes associated with monocyte-derived DCs, including Cd209a, Cd74, and Nr4a3 (Boulet et al., 2019; Ponichtera et al., 2014; Stables et al., 2011) (Figure 2F and Supplemental Table 2C). The remaining non-DC MHCII^-^ fraction was enriched for genes that are known to regulatory function in inflammation, lipid metabolism and angiogenesis including Crip2, Fxyd2, and Rnase4 (Cheung et al., 2011; Li et al., 2013; Mayan et al., 2018) (Figure 2G and Supplemental Table 2C). Thus, we conclude that the MHCII^-^ compartment represents a pure population of tissue-resident synovial Ly6C^-^ cells.

We compared our single-cell data to those recently published using mouse and human synovium. The differentially expressed genes associated with subpopulations 2, 4 and 5 (Supplemental Table 2A) as well as MHCII^+^ and MHCII^-^ cells (Supplemental Table 2C) were compared to the top 20 marker genes for the single-cell myeloid subpopulations (excluding neutrophils) identified by Culemann et al. at day 5 post-STIA (Culemann et al., 2019) (Figure 2H, Supp Table 2D). Consistent with our annotations, subpopulations 2 and 4 demonstrate high unique overlap with STMN1+ proliferating cells and CCR2+IL1B+ infiltrating macrophages, respectively using data from Culemann et al (p<0.001, FWER=0.05). In contrast, the MHCII^-^ cells appear to be split between Relmα interstitial myeloid cells (p=0.0009) and CCR2^+^Arg^+^ infiltrating myeloid cells (p=2.65×10^-8^), while MHCII^+^ overlapped with MHCII^+^ macrophages (p=5.89×10^-6^) and MHCII^high^ DCs (p<1.75×10^-8^) from Culemann et al. These results suggest that our subpopulations based on enriching for CD64-Tim4-cells do not exactly match the Culemann clusters. Next, using published data from the AMP consortium on CD14^+^ myeloid cells sorted from human synovium (Zhang et al., 2019), we annotated cells with high expression of genes associated with our murine MHCII^+^ and MHCII^-^ profiles (Figures 2I, S2M-N and Supplemental Table 2C). We found that the proportion of MHCII+ to MHCII-cells was highest in AMP patients annotated as Leukocyte-rich RA compared with Leukocyte-poor RA or OA patients. Similarly, the MHCII+ cells were more likely to be annotated in the AMP IL1B+ and C1QA+ clusters that were associated with active RA and this was supported by a significant overlap in marker genes (p<0.0025, FWER=0.05). (Figures 2J, S2O-P, and Supplemental Table 2D). We also examined the overlap between genes implicated in our subpopulations and markers for 9 newly defined subpopulations of RA synovial macrophages using scRNA-seq from Alivernini et al (Alivernini et al., 2020). Our MHCII^-^ cells exhibited significant association with Alivernini CD48^high^SPP1^pos^ cells (p=2.56×10^-5^), while MHCII^+^ cells were associated with both the FOLR2^+^ID2^+^ and FOLR2^+^ICAM1^+^ cells (p<0.001, FWER=0.05) (Figure S2Q, Supplemental Table 2D).

We then performed scRNA-seq on CD45^+^CD11b^+^Ly6G^-^SigF^-^Tim4^-^CD64^-^ synovial cells from NR4A1^-/-^ and CCR2^-/-^ mice to determine whether their compositions were affected by CCR2 or NR4A1 deficiency (Figure 2K, Supplemental Table 2E). These datasets were integrated with control cells (C57Bl/6) and the subpopulation annotation was superimposed (Figure S2R). The distribution of CCR2^-/-^ and NR4A1^-/-^cells across the 6 subpopulations was significantly different from controls (p<2.2e-16, p=4.89e-10) (Figure 2L). Although there was no significant difference in the proportion of cells annotated as MHCII^-^ vs. MHCII^+^ in CCR2^-/-^ mice, we observed an increase in the proportion of MHCII-cells in NR4A1^-/-^ mice (Figure 2M). Collectively, these data demonstrate that MHCII^-^ cells represent a distinct Ly6C^-^ population in the synovium of mice and RA patients, which are independent of CCR2 and NR4A1.

### Synovial Ly6C^-^ cells exist as two separate populations

To better define the location of Syn Ly6C^-^ cells, we used an *in vivo* intra-vascular labeling system (Jakubzick et al., 2013). Only intra-vascular cells were labelled by administering intravenously (I.V.) a fluorescently conjugated anti-CD45 antibody (αCD45-BUV661-(I.V.) prior to euthanasia (Figure 3A). Over 90% of the circulating leukocytes were labeled 5 min post I.V. (Figure S3A). Thus, immune cells co-labeled with αCD45-BUV661-(I.V.) and the *ex vivo* (E.V.) CD45 antibody (αCD45-AF700-(E.V.) were intra-vascular, while single positive αCD45-AF700-(E.V.) cells were extra-vascular. Synovial macrophages were distinguished using CD64^+^ and Tim4^+^ and confirmed by positive staining of CD204 and CD63 (Figures 3B, S3B). Intra-and extra-vascular cells were then gated based on CD45^+^CD11b^+^Ly6G^-^Tim4^-^CD64^-^Ly6C^-^ expression to isolate the synovial Ly6C^-^ cells. Intra-vascular synovial Ly6C^-^ cells (i.v. Syn Ly6C^-^) were gated for cell surface expression of CX3CR1 and CD43 (Figures 3B, S3C-D). Extra-vascular (e.v.) cells were divided into e.v. Syn Ly6C^-^ and mono-DC based on expression of MHCII, with mono-DC expressing significantly higher levels of MHCII than the other cell populations (p<0.001) (Figures 3B, S3C-D), consistent with the scRNA-seq data (Figure 2). The e.v. Syn Ly6C^-^ cells were the largest population, with a mean 26,075 ±4608 cells, compared to i.v. Syn Ly6C^-^ cells (355 ±64) and mono-DCs (9,851 ±1660) (Figure 3C). The inconsistency with the scRNA-seq findings where we defined a much smaller MHCII^-^ populations is indicative of our very conservative cutoff for MHCII expression and the difference between gene expression (RNA-seq) as opposed to surface marker expression of the protein (flow cytometry).

**Figure 3:**
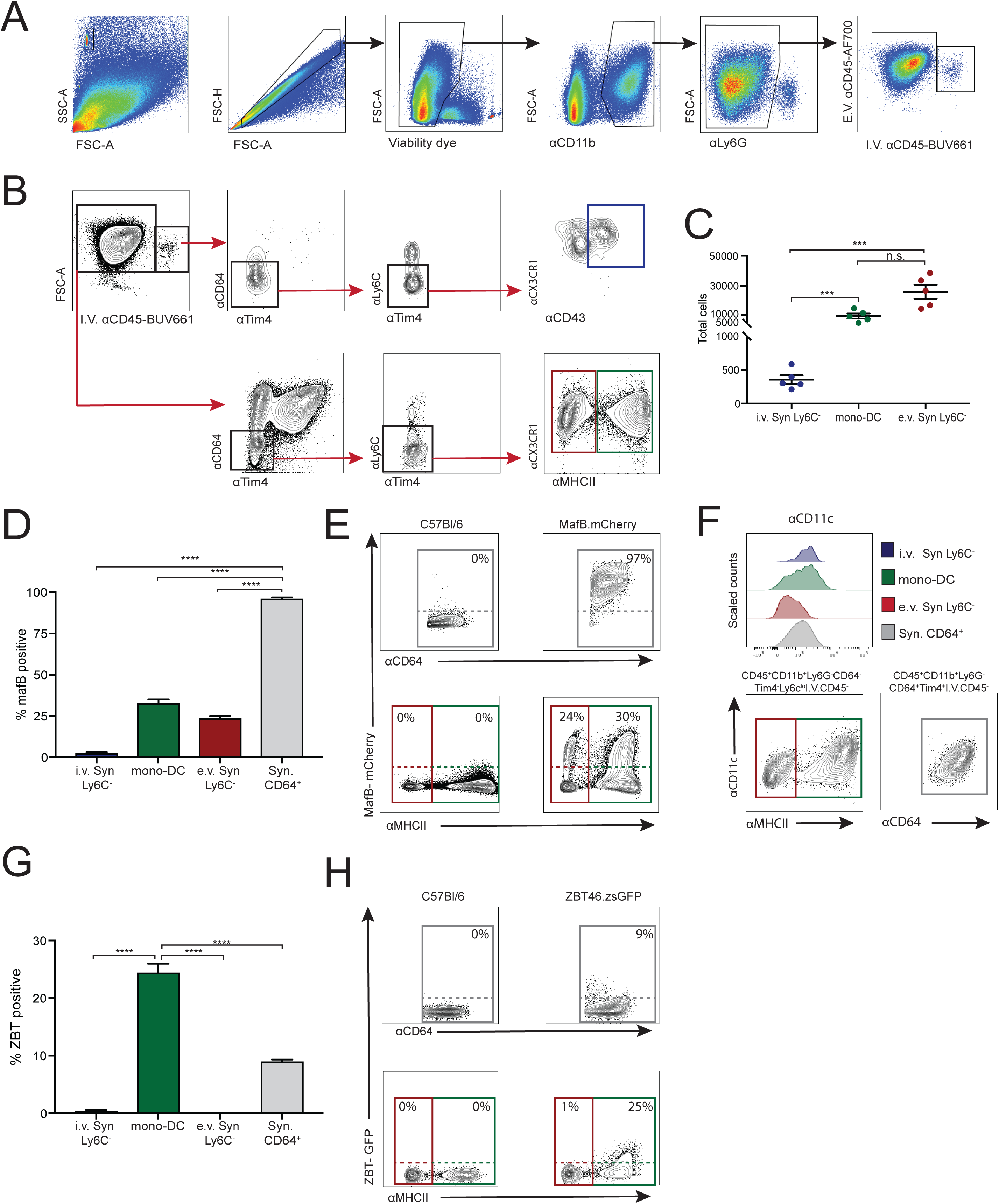
Identification of intra- and extra-vascular Syn Ly6C^-^ cells by flow cytometry. A) Gating strategy to distinguish adherent intra-vascular and extra-vascular myeloid cells and B) intra- and extra-vascular NCM and mono-DC. C) Numbers of NCM and mono-DC in hindjoints of C57Bl/6 mice in steady state. D-E) Expression of MafB-mcherry, F) CD11c, G-H) ZBT46 in Syn Ly6C^-^ cells, mono-DC, and synovial CD64^+^ cells. All graphs are mean N>4 ±SEM P-value was calculated using unpaired t-test. ***=p<0.005, ****=p<0.001.

MafB is a known transcription factor (Wu et al, 2016) required for macrophage development. We utilized MafB.mcherry knock-in reporter mice to measure MafB expression in the synovium. The i.v. Syn Ly6C^-^ cells (2.9 ±0.4%, p=0.004), mono-DC (33.2±1.8%, p=0.006) and e.v. Syn Ly6C^-^ cells (23.9 ±1.2%, p=1.9 ×10^-9^) were markedly less positive for mCherry compared to synovial macrophages (96 ±0.5%) (Figure 3D-E). PB NCM had no detectable expression (Figure S3E).

We then examined cell surface expression of CD11c as well as measured expression of the DC gene Zbt46 using zDC-cre mice crossed with zsGFP reporter mice (zDC-GFP). i.v. Syn Ly6C^-^ cells and mono-DCS expressed higher levels of CD11c than e.v. Syn Ly6C^-^ cells (p<0.05) (Figures 3F, S3D), consistent with the known pattern of CD11c on monocyte-derived cells and DCs. In the zDC-GFP mice, expression of GFP was most widespread in mono-DCs with 24.5 ±1.5% GFP^+^ cells, but was not significantly expressed in either Syn Ly6C^-^ populations or in PB monocytes (Figures 3G-H, S3E). Thus, these data uncover two central Ly6C^-^ populations in the synovium that can be distinguished by their location relative to the blood vessels, i.e. intra- and extravascular.

### Intra-and extra-vascular synovial Ly6C^-^ exhibit different functionality

To investigate the properties of the two synovial Syn Ly6C^-^ cells and contrast them with mono-DCs, we compared the transcriptional profiles of these cells using bulk RNA-seq. Each population was highly reproducible across replicates and characterized by unique transcriptional profiles (Figures 4A, S4A). We further confirmed the relationship of the bulk-sorted cells to the subpopulations identified by scRNA-seq using singleR with the bulk data as reference. The majority of cells assigned to i.v. Syn Ly6C^-^ cells were from scRNA-seq subpopulation 4 (Figures 4B, S4B-C). In contrast, cells assigned to mono-DC and e.v. Syn Ly6C^-^ cells largely comprised scRNA-seq MHCII^+^ and MHCII^-^ cells, respectively (Figures 4B, S4D-E). i.v. Syn Ly6C^-^ cells expressed the highest levels of Cx3cr, Spn (CD43), Cebpb and Nr4a1, genes associated with PB NCM (Figure 4C). Further, the mono-DC population identified by flow cytometry shared gene expression (Figure 4D) with scRNA-seq MHCII^+^ cells (Figure S4C) including Cd74, Cd209a, Zbtb46 and Flt3. Meanwhile, e.v. Syn Ly6C^-^ cells were enriched for genes expressed in scRNA-seq MHCII^-^ cells (Figure S4C) such as Alox5, CD177, Fxyd2 and Rnase4 (Figure 4E).

**Figure 4:**
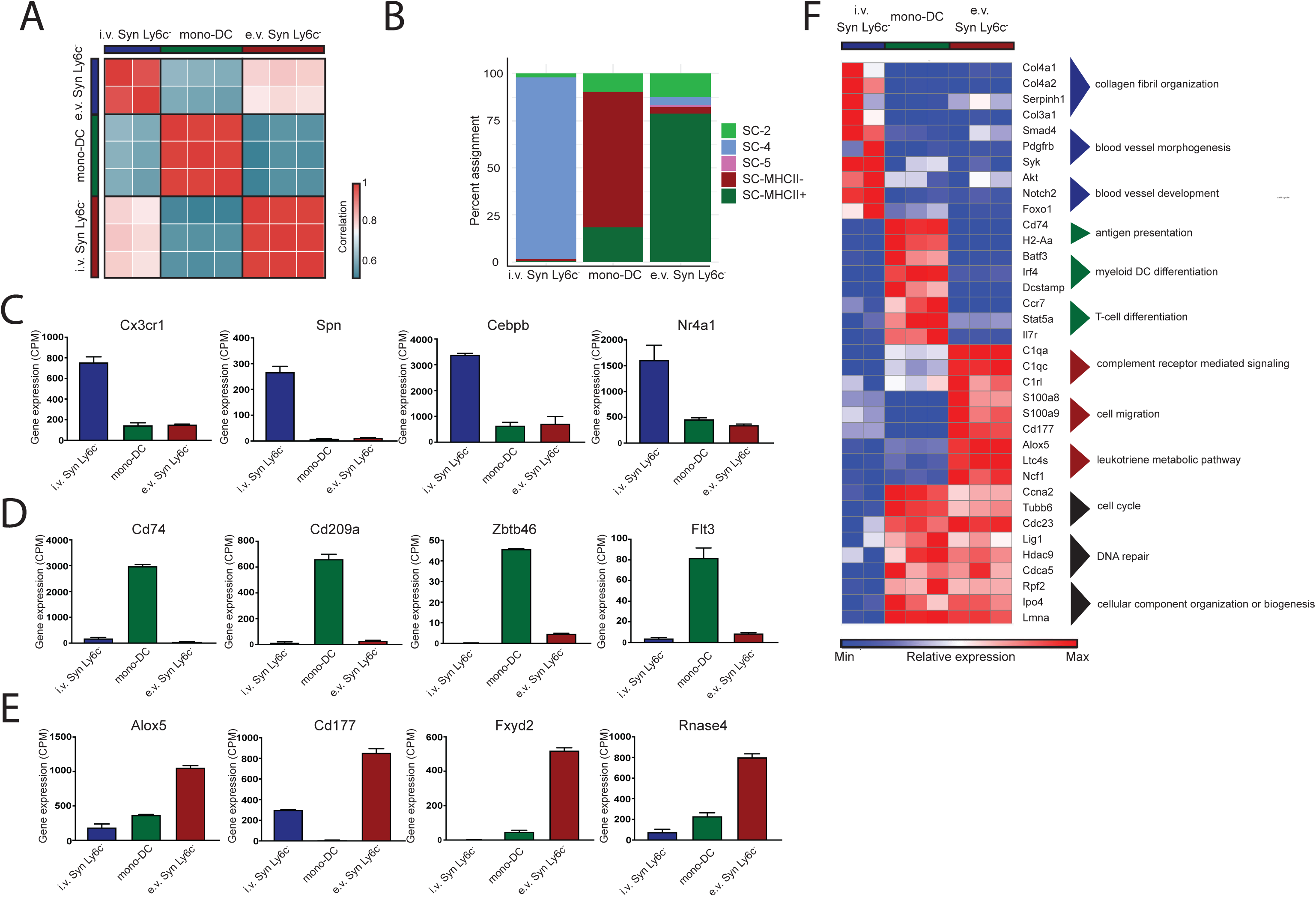
Extra-vascular tissue location confers phenotype of Syn Ly6C^-^ cells. A) Pairwise Pearson’s correlation of global gene expression between replicates of i.v. Syn Ly6C^-^, mono-DC and e.v. Syn Ly6C^-^ cells. B) Percent of cells assigned to i.v. Syn Ly6C^-^, mono-DC or e.v. Syn Ly6C^-^ cells from each scRNA-seq subpopulation or MHCII^+/-^ compartment (Figure 2) using the bulk transcriptional data as reference in singleR. Genes with preferential expression in C) i.v.NCM, D) mono-DC and E) e.v. Syn Ly6C^-^. 2. F) Relative expression of representative genes from GO processes associated with i.v. Syn Ly6C^-^ cells, mono-DC, e.v. Syn Ly6C^-^ cells.

We then performed k-means clustering on 5127 differentially expressed genes across these populations to define 4 clusters: one each with expression specific to i.v. Syn Ly6C^-^ cells (1), mono-DC (2) and e.v. Syn Ly6C^-^ cells (3) and a final cluster (4) preferentially expressed in both mono-DC and e.v. Syn Ly6C^-^ cells (Figure S4D, Supplemental Table 3A). The i.v. Syn Ly6C^-^ cell-specific cluster was enriched for genes associated with collagen fibril organization (Col6a1/2, Col4a1/2, Adamts2), blood vessel morphogenesis (Smad4, Pdgfra, Syk). and blood vessel development (Akt1, Notch2, Foxo1) (Figure 4F, Supplemental Table 3B). As expected, the mono-DC-specific cluster was enriched for genes involved in MHCII antigen presentation molecules (H2-Aa, Cd74), as well as genes involved in myeloid DC differentiation (Irf4, Dcstamp, Btf3), and genes that play a role in T-cell differentiation (Ccr7, Stat5a, Tnfsf9) (Figure 4F and Supplemental Table 3C). For the e.v. Syn Ly6C^-^ cell-specific cluster, enriched GO processes included complement receptor mediated signaling (C1qa/c, C5ar1, Fcna/b), and chemotaxis, cell motility, locomotion, and cell migration pathways that overlap with S100a8/9, Elane and Cd177 genes (Figure 4F and Supplemental Table 3D). Additionally, the leukotriene metabolic pathway was enriched in this cluster with genes including Alox5, Ltc4s, and Ncf1. The cluster shared between mono-DC and e.v. Syn Ly6C^-^ included cell cycle (Ccna2, Tubb6, Cdc23), DNA repair (Lig1, Hdac9, Cdca5) and cellular component organization or biogenesis (Rpf2, Ipo4, Lmna) (Figures 4F, S4E, Supplemental Table 3E).

We then examined cell surface expression of receptors that were associated with transcriptional profile of Syn Ly6C^-^ cells, mono-DC and Syn CD64^+^ cells (Figure S4F). FcγRIV and Treml4 separated the i.v. Syn Ly6C^-^ cells from the e.v. Syn Ly6C^-^ cells as well as mono-DC and Syn CD64^+^ cells, while CD177 discriminates the e.v. synovial from the i.v. Syn Ly6C^-^ cells, mono-DC and Syn CD64^+^ cells. Although the gene expression of CD88, Folrb and Lyve1 was highest in e.v. Syn Ly6C^-^ cells as compared to i.v. Syn Ly6C^-^ cells or mono-DC, Syn CD64^+^ cells displayed the greatest intensity of staining.

### e.v. Syn Ly6C^-^ cells are long lived with embryonic origins

Tissue-resident macrophage populations have been shown to originate from either the fetal liver or adult hematopoiesis, but the origins of tissue Syn Ly6C^-^ cells are unknown. We crossed a tamoxifen-inducible CX3CR1-Cre (CX3CR1^CreER^) mouse with a GFP reporter mouse (zsGFP) to generate a mouse (CX3CR1^CreER^.zsGFP) in which CX3CR1^+^ cells express GFP after administration of tamoxifen (TMX). Mice were treated with 2 doses of TMX over subsequent days to measure the steady-state lifespan of circulating and synovial cells. The vast majority of PB monocytes were readily positive for GFP, which was lost by day 14 (Figure 5A-B), in line with the previously reported lifespan of PB NCM (Yona et al., 2013). In contrast, 18 ±0.5% of the Syn Ly6C^-^ cells were GFP^+^ at D3 but this level remained stable throughout the 21-day time course (Figure 5B-C). Syn CD64^+^ cells also displayed 24 ±0.9% GFP+ over the 21 days (Figure 5B-C), consistent with previous studies suggesting that this population does not proliferate (Culemann et al., 2019; Misharin et al., 2014). These data suggest that the e.v Syn Ly6C^-^ cells are long lived, distinct from short-lived circulating monocytes and do not differentiate into synovial macrophages.

**Figure 5:**
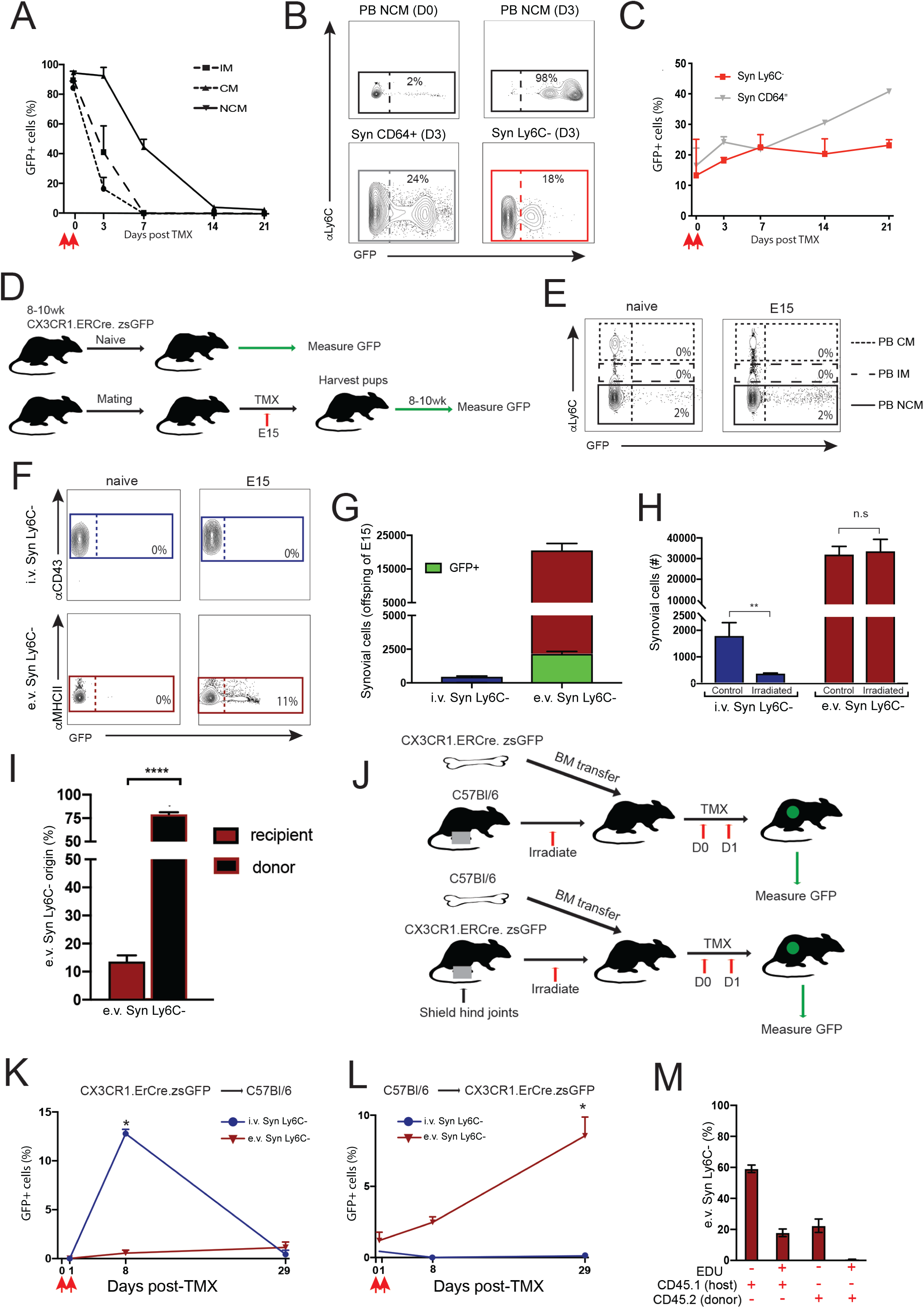
e.v. Syn Ly6C^-^ cells are fetal liver derived and long lived. A) GFP expression in PM monocytes, B) CD64^+^, and C) Syn Ly6C^-^ cells from adult CX3CR1^Cre.ER^.zsGFP mice treated with 50mg/kg TMX at D0 analyzed on D1. D) Experimental approach for lineage tracing. E) Labeling of PB monocytes and F-G) I.v. and e.v. Syn Ly6C^-^ cells in adult CX3CR1^Cre.ER^.zsGFP without TMX, and offspring of CX3CR1^Cre.ER^.zsGFP mice treated with TMX on day 15 of gestation (E15). H) Total i.v. and e.v. Syn Ly6C^-^ cells in steady-state and 24hrs post-non-lethal irradiation. I) Percentage e.v. Syn Ly6C^-^ cells at 8 weeks post-reconstitution in irradiated CD45.1-> CD45.2 bone marrow chimeras. J) Experimental approach for shielded chimeras. K) Percentage of GFP^+^ i.v. and e.v. Syn Ly6C^-^ cells in the synovium in CX3CR1^Cre.ER^.zsGFP -> C57Bl/6 bone marrow chimeric mice and L) C57Bl/6 -> CX3CR1^Cre.ER^.zsGFP 7 bone marrow chimeras and 28 days after administration of 50mg/kg TMX. M) Edu expression in e.v. Syn Ly6C^-^ cells after 14 days post-EdU treatment in CD45.2-> CD45.1 bone marrow chimeric mice. All graphs are mean N>4 +SEM p-value was calculated using unpaired t-test. *=p<0.05, **=p<0.01, ****=p<0.001.

We then utilized CX3CR1^CreER^.zsGFP mice for Syn Ly6C^-^ cell fate-mapping studies (Figure 5D). TMX was delivered at E15 to pregnant mothers from the cross of CX3CR1^CreER^ with zsGFP mice, which will allow for identification of embryonically derived cells that are not derived from the yolk sac. TMX-naïve CX3CR1^CreER^.zsGFP mice provided control to estimate background levels of GFP. PB monocytes from progeny of TMX-treated E15 mice displayed undetectable levels of GFP expression similar to background in TMX-naïve mice (Figure 5E). Meanwhile 11 ±1.5% of e.v. Syn Ly6C^-^ cells were GFP^+^ in the progeny of the E15 TMX-treated CX3CR1^CreER^ X zsGFP pregnant females compared to 0% in i.v. Syn Ly6C^-^ cells (Figure 5F-G). Neutrophils (Ly6G^+^) were also negative for GFP, while the majority of synovial macrophages (66 ±3%) were GFP^+^ (Figure S5A-B). We also subjected C57Bl/6 mice to non-lethal gamma irradiation-induced cell death to further support the tissue-residency of e.v. Syn Ly6C^-^ cells. We found that the bulk of e.v. Syn Ly6C^-^ cells but not i.v. Syn Ly6C^-^ cells were initially refractory towards non-lethal gamma irradiation at one day post irradiation (Figure 5H). Further, a population of the host e.v. Syn Ly6C^-^ cells was detectable in CD45.1/CD45.2 BMC mice at 8 weeks post-transplant (Figure 5I). These data support that even when the synovial niche is disrupted a subpopulation of e.v. Syn Ly6C^-^ cells are radio-resistant and may self-replenish.

Given that e.v. Syn Ly6C^-^ cells may originate from the fetal liver and that a subpopulation is radio-resistant, we examined whether the e.v. Syn Ly6C^-^ compartment was replenished by PB monocytes over time. C57Bl/6 mice were treated with non-lethal gamma-irradiation but had their hind joints shielded to retain synovial compartment and received CX3CR1^CreER^zsGFP bone marrow (CX3CR1^CreER^zsGFP->C57Bl/6) immediately after irradiation. TMX was administered on experimental D0 and D1 at 8 weeks post-irradiation and transfer. GFP was analyzed directly after the second dose of TMX (D1), and at 7 and 28 days post-TMX (Figure 5J). PB monocytes from CX3CR1^CreER^zsGFP->C57Bl/6 chimeric mice displayed a dramatic increase in GFP expression, which eventually disappeared by 28 days post-TMX administration (Figure S5C). Over 12% of the i.v. Syn Ly6C^-^ cells were GFP^+^ in the CX3CR1^CreER^zsGFP->C57Bl/6 chimeric mice on D7 post-TMX, which was reduced to undetectable levels by day 28 post-TMX. These data support i.v. Syn Ly6C^-^ cells are derived from donor PB monocytes and are short-lived (Figure 5K). In contrast, the expression of GFP was not significantly above background in e.v. Syn Ly6C^-^ cells throughout the 28-day time course, which indicates that the donor monocytes do not contribute to e.v. Syn Ly6C^-^ cells when the niche is not disrupted. Furthermore, a reverse chimera was also generated (C57Bl/6->CX3CR1^CreER^zsGFP) (Figure 5J) and the hind joints were protected by lead shielding of host mice. In these chimeric mice, the i.v. Syn Ly6C^-^ cells were negative for GFP throughout the 28 days post-TMX further supporting that the i.v. Syn Ly6C^-^ cells are derived from circulating monocytes. In contrast, only a small fraction of e.v. Syn Ly6C^-^ cells (2.5 ±0.3%) were GFP^+^ cells at D7, which increased to 8.5 ±1.3 by Day 28 (p=0.025) (Figure 5L). Since the numbers of GFP^+^ e.v. Syn Ly6C^-^ cells are reduced in C57Bl/6->CX3CR1^CreER^zsGFP chimeric mice as compared to CX3CR1^CreER^zsGFP mice at steady state (Figure 5C), these data suggest that e.v. Syn Ly6C^-^ niche is retained locally. To address whether the increase in GFP^+^ e.v. Syn Ly6C^-^ cells over time in C57Bl/6->CX3CR1^CreER^zsGFP chimeric mice support the self-replenishment of e.v. Syn Ly6C^-^ cells, we continuously administered nucleotide analog EdU for 14d to 8-week post-irradiated and hind joint-shielded CD45.2 CD45.1 bone marrow chimeric mice. Over 17.8±1.4% of CD45.1 (host) e.v. Syn Ly6C^-^ cells were EdU^+^ after 14 days, whereas <1% of CD45.2 donors e.v. Syn Ly6C^-^ cells were EdU^+^ (Figure 5M). Collectively, these data suggest the e.v. Syn Ly6C^-^ cells are derived embryonically, radioresistant, and capable of self-renewal but can be replenished from circulating monocytes when the synovial niche is disrupted.

### Rapid expansion of e.v. Syn Ly6C^-^ cells in the synovium is required for development of inflammatory arthritis

To assess the dynamics of e.v. Syn Ly6C^-^ cells during induction of inflammatory arthritis, we determined the numbers of Syn Ly6C^-^ cells at 1 and 24 hours post-STIA. i.v. and e.v. Syn Ly6C^-^ cells increased by 4- (p=0.009) and 2-fold (p=0.008), respectively at 1h compared to steady state, but only e.v. Syn Ly6C^-^ cell numbers continued to expand (2-fold) from 1 to 24 hours (Figure 6A, Figure S6A). In contrast, neutrophils in the synovium were not significantly changed during the first 24 hours post-STIA, consistent with previous studies (Misharin et al., 2014). These results demonstrate that Syn Ly6C^-^ expansion precedes neutrophil infiltration.

**Figure 6:**
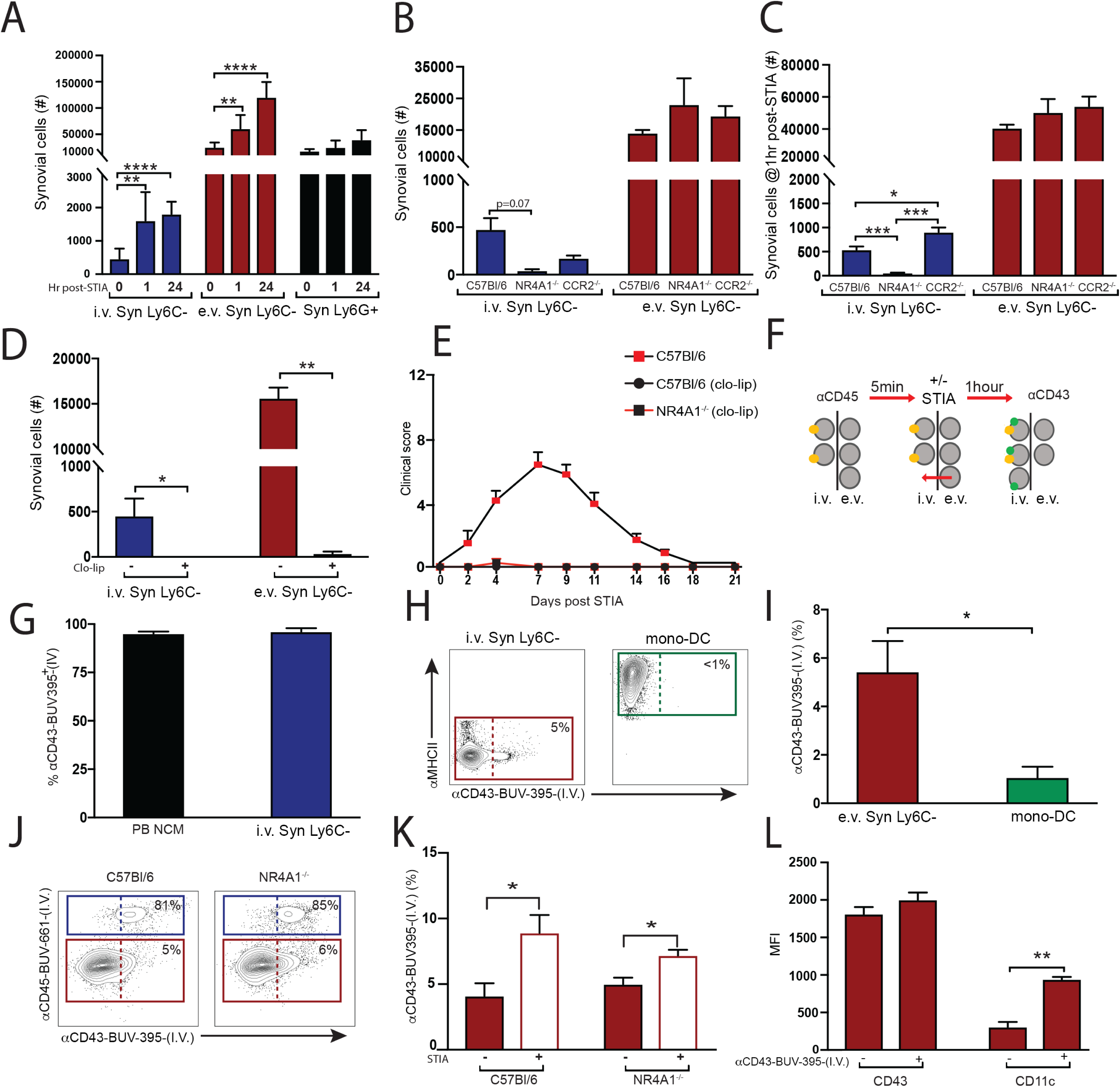
Extra-vascular Syn Ly6C^-^ cells are the critical population for development of inflammatory arthritis. A) Numbers of i.v. and e.v. Syn Ly6C^-^ cells and Syn Ly6G^+^ cells at 1 and 24hrs post STIA. B) Numbers of i.v. and e.v. Syn Ly6C^-^ cells in steady state and C) at 1h post-STIA in C57Bl/6, NR4A1^-/-^ and CCR2^-/-^ mice. D) Depletion of i.v. Syn Ly6C^-^ cells and e.v. Syn Ly6C^-^ cells post treatment with 200μL of Clo-lip. E) Clinical scoring of STIA induced 24h after Clo-lip treatment in C57Bl/6 and NR4A1^-/-^ mice. F) Experimental design of reverse-extravasation system. G) Labeling of PB and i.v. Syn Ly6C^-^ cells via IV administration of anti-CD43 antibody H-I) Labeling of e.v. Syn Ly6C^-^ cells and mono-DC with I.V. αCD43-BUV395 after 1h steady state. J) Labeling of e.v. Syn Ly6C^-^ cells and i.v. Syn Ly6C^-^ cells with anti-CD43 antibody in C57Bl/6 and NR4A1^-/-^ after 1h steady state and K) 1h after STIA. L) Surface expression of CD43 and CD11c on I.V. αCD43-BUV395^+/-^ e.v. Syn Ly6C_-_ cells. Graphs are mean N>4 +SEM P-value was calculated with unpaired t-test. *=p<0.05, **=p<0.01, ***=p<0.005, ****=p<0.001.

We then examined the e.v. Syn Ly6C^-^ cell response in NR4A1^-/-^ and CCR2^-/-^ mice to determine whether the expansion of e.v. Syn Ly6C^-^ cells requires contribution from PB NCM or PB CM. Under steady state conditions NR4A1^-/-^ and CCR2^-/-^ mice have reduced numbers of i.v. Syn Ly6C^-^ cells, and comparable levels of e.v. Syn Ly6C^-^ cells to C57Bl/6 mice (Figure 6B). One hour after administration of STIA, i.v. Syn Ly6C^-^ cells numbers in NR4A1^-/-^ mice remained lower than C57Bl/6 (p=0.0003), and CCR2^-/-^ (p=0.0001), which underwent even more significant expansion than C57Bl/6 (p=0.02) (Figures 6C, S6B). Meanwhile e.v. Syn Ly6C^-^ cells levels remained comparable across genotypes, having undergone similar levels of expansion 1h post-STIA (Figure 6C, Figure S6B). To determine if e.v. Syn Ly6C^-^ cells are a critical population in STIA, we examined whether clodronate-laden liposomes (Clo-lip) would deplete i.v. and e.v. Syn Ly6C^-^ cells. Both i.v. and e.v. Syn Ly6C^-^ cells are markedly reduced in response to Clo-lip at 24 hrs (p= 0.007, p=0.006) (Figure 6D). As expected Clo-lip treatment depleted circulating monocytes but did not affect the numbers of Syn CD64+ cells (Figure S6C-D), similar to previous studies (Misharin et al., 2014). Administration of Clo-lip prior to STIA completely prevented disease in both C57Bl/6 and NR4A1^-/-^ mice (Figure 6E). Given the redundancy of i.v. Syn Ly6C^-^ cells in NR4A1^-/-^ mice, these data suggest that the e.v. Syn Ly6C^-^ cells are essential for inflammatory arthritis.

### Extravascular Syn Ly6C^-^ cells are capable of reverse extravasation

We observed that e.v. Syn Ly6C^-^ cells are sensitive to Clo-lip while synovial macrophages are unaffected. These results may be explained if e.v. Syn Ly6C^-^ cells have access to the circulation. To investigate this notion, we modified the *in vivo* labeling system to include a second I.V. antibody (Figure 6F), αCD43-BUV395-(IV), which labels 95 ±1.1 % of circulating NCM and 96 ±1.8% of i.v. NCM (Figures 6G, S6E). With this system of administering αCD45-BUV661-(IV) 1h prior to αCD43-BUV395-(IV), i.v. Syn Ly6C^-^ cells that are retained in the intra-vascular space remain dual positive for I.V. αCD45-BUV661 and I.V. αCD43-BUV395, while e.v. Syn Ly6C^-^ cells that transverse to circulation within the hour will only be positive for αCD43-BUV395-(IV) (Figure 6H). Using this strategy, 5 ±1.2% of e.v. Syn Ly6C^-^ cells labeled with αCD43-BUV395-(IV) during steady state, whereas only 1 ±0.4% of mono-DC became labeled, confirming that a fraction of e.v. Syn Ly6C^-^ cells undergo reverse extravasation during a 1hour duration (Figure 6H-I). A similar trend was observed in NR4A1^-/-^ mice (Figure 6J-K). Further, the addition of STIA as an inflammatory stimulus increased the reverse transmigration by 2-fold of e.v. Syn Ly6C^-^ cells, indicative by the positive staining for αCD43-BUV395-(IV) in C57Bl/6 (p=0.045) and NR4A1^-/-^ mice (p=0.025) within 1h post-STIA. Since e.v. Syn Ly6C^-^ cells undergo enhanced reverse extravasation in response to STIA, these data suggest that loss of e.v. Syn Ly6C^-^ cells are exposed to the vasculature.

Previous studies have shown monocytes in liver undergo reverse extravasation associated with enhanced expression of CD11c (Zimmermann et al., 2016). Similarly, the e.v. Syn Ly6Ccells that underwent reverse extravasation i.e. αCD43-BUV395^+^-(IV) exhibited a significant elevation in CD11c expression (p=0.007) (Figure 6L), compared to the cells that remained extravascular, whereas the level of CD43 remained consistent (Figure 6L, Figure S6F). Together, these data and the Clo-lip studies demonstrate that e.v. Syn Ly6C^-^ cells undergo reverse transmigration and have continued access to the vasculature.

### LFA1 is required for STIA induced reverse extravasation of e.v. Syn Ly6C^-^ cells

Previously studies have shown that LFA is required for leukocyte extravasation into tissue and for STIA (Watts et al., 2005). LFA1^-/-^ mice failed to develop inflammatory arthritis (Figure 7A) consistent with other’s work (Monach et al., 2010; Watts et al., 2005). There were no significant differences in the numbers of PB CM, PB NCM, i.v. and e.v. Syn Ly6C^-^ cells between LFA1^-/-^ mice and control mice at steady state (Figure 7B-D). However, neither LFA^-/-^ Syn Ly6Cpopulation exhibited the expansion in response to arthritogenic serum (Figure 7E) observed in C57Bl/6 mice (Figure 6A). In addition, LFA1^-/-^ e.v. Syn Ly6C^-^ cells did not undergo increased reverse extravasation (measured with αCD43-BUV395-(IV)) 1hr following STIA as compared to C57Bl/6 mice (Figure 7F).

**Figure 7:**
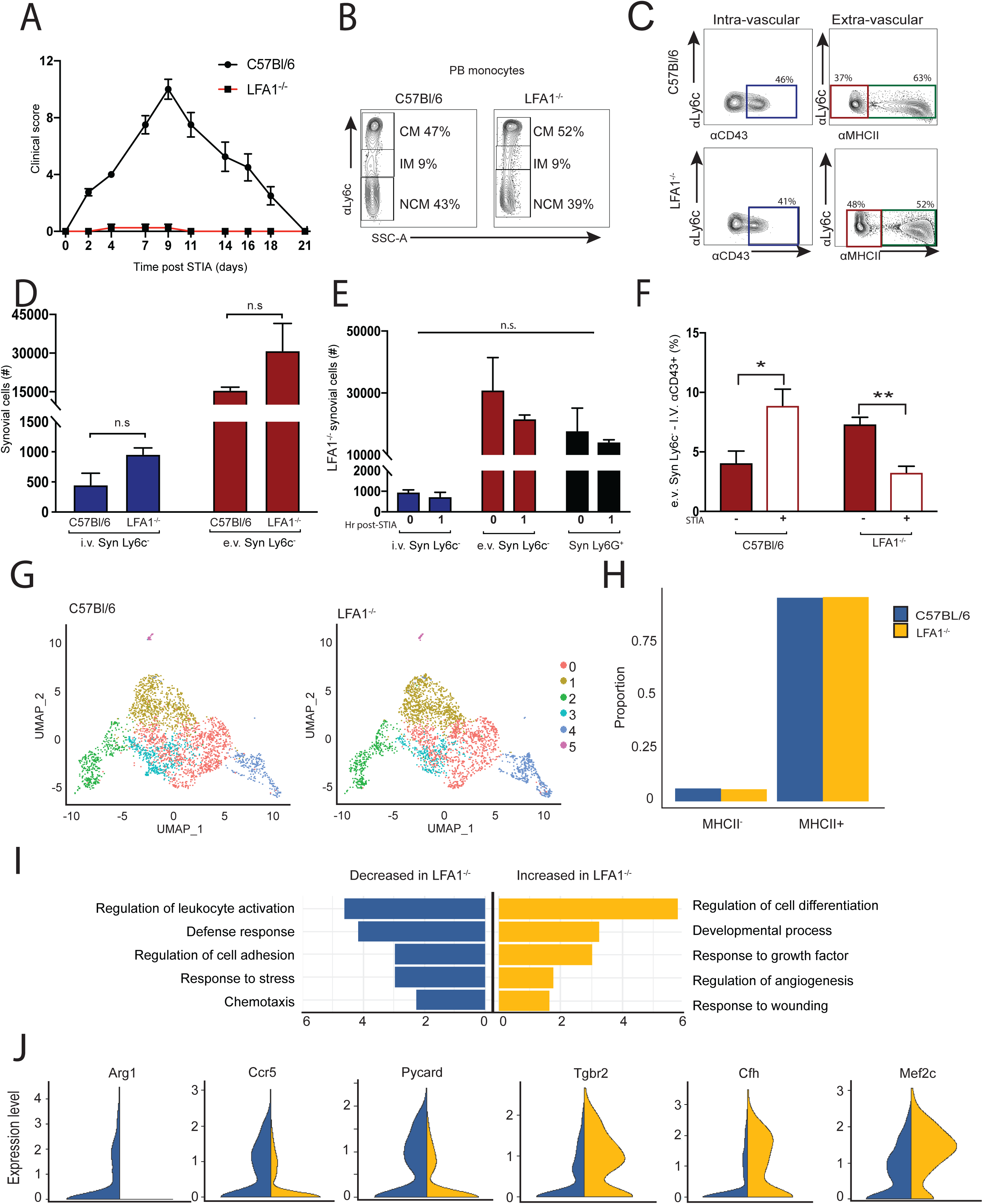
Deletion of LFA1 reduced pro-inflammatory phenotype of e.v. Syn Ly6C^-^ cells. A) STIA clinical score in C57Bl/6 and LFA1^-/-^ mice. B) PB monocytes and C-D) i.v. Syn Ly6C^-^, and e.v. Syn Ly6C^-^ cells in C57Bl/6 and LFA1^-/-^ in steady-state. E) Number of i.v. Syn Ly6C^-^, e.v. Syn Ly6C^-^ cells and Syn Ly6G^+^ cells 1hr post-STIA in C57Bl/6 and LFA1^-/-^ mice. F) e.v. Syn Ly6C^-^ cells labeled with I.V αCD43-BUV395 in steady state and 1hr post-STIA. G) Integration of scRNA-seq data on CD45^+^CD11b^+^Tim4^-^CD64^-^ cells from LFA^-/-^ mice with C57Bl/6. To obtain comparable numbers, both datasets were sub-sampled to 3000 cells. H) Ratio of cells annotated as either MHCII^+^ or MHCII^-^ in C57Bl/6 and LFA1^-/-^ mice. I) Selected GO processes associated with differentially expressed genes in the MHCII^-^ compartment (representing e.v. Syn Ly6C^-^ cells) between LFA1^-/-^ and C57Bl/6 mice. J) Ridge plots of representative genes that are increased or decreased in expression in MHCII^-^ cells from LFA1^-/-^ compared with C57Bl/6 mice. Graphs are mean N>4 +SEM P-value was calculated with unpaired t-test. *=p<0.05, **=p<0.01.

### LFA1 deletion reduces pro-inflammatory transcriptional profile of e.v. Syn Ly6C^-^ cells

To determine how LFA1 deletion affected Syn Ly6C^-^ cells on the transcriptional level, we performed scRNA-seq on isolated CD45^+^CD11b^+^Ly6G^-^SigF^-^Tim4^-^CD64^-^ synovial cells from LFA1^-/-^ mice at steady state. As before, we integrated and superimposed the 6 subpopulations (0-5) defined from C57Bl/6 mice on 3500 LFA1^-/-^ cells (Figures 7G, S7A, Supplemental. Table 4C). LFA1^-/-^ mice displayed an altered distribution of cells across the 6 subpopulations (p<2.2×10^-16^) (Figure S7B,C) but the ratio of MHCII^+^ to MHCII^-^ cells was comparable to C57Bl/6 mice (Figure 7H). LFA^-/-^ e.v. Syn Ly6C^-^ cells exhibit decreased expression of genes associated with chemotaxis (Ccl17, Itgb2, Ccl9), defense response (Arg1, Mif, Cfp, Itgax), regulation of cell adhesion (Ccr5, Lgals3, Adam8) and stress response (Pycard, Flt1, Prdx5, Vegfa) (Figure 7I-J, Supplemental Table 4A-B). In contrast, genes associated with regulation of cell differentiation (Fos, Cd36, Mef2c, Mafb, Csf1r), regulation of angiogenesis (Tcf4, Pf4, Tgfbr2), and response to wounding (Macf1, Aqp1, Cfh) were increased in expression in LFA^-/-^ e.v. Syn Ly6C^-^ cells compared to C57Bl/6 (Figure 7I-J, Supplemental. Table 4A-B). Based on these data, we demonstrate that e.v. Syn Ly6C^-^ cells play a vital role in suppressing inflammation by controlling the chemotaxis of leukocytes such as granulocytes to the synovium.

## Discussion

Over the past several years, numerous studies have characterized CM and NCM in circulation and contrasted these with differentiated macrophages in the tissue. Our study is the first to identify two synovial populations that exist in the tissue but do not exhibit canonical macrophage markers. First, we described a distinct population of i.v. Ly6c^-^ cells, that are transcriptionally similar to PB NCM and require NR4A1 but remain attached to the vessel wall even after perfusion. Moreover, we identified another e.v. synovial Ly6C^-^ population that is negative for CD64 and Tim4 and is transcriptionally distinct from both circulating monocytes and synovial macrophages. These Syn. Ly6C^-^ cells are long-lived, embryonically derived terminal cells that are self-proliferative and do not require CCR2, NR4A1 or LFA for development. A recent study described a population of Ly6c^lo^ cells localized to the e.v. tissue of the lung that they termed monocytes (Schyns et al, 2019). However, these lung cells were derived from circulating NCM, expressed CD64, and provided a niche to replace lung interstitial macrophages, suggesting they are in a transitional state between monocytes and macrophages.

Prior studies on the role of monocytes in RA have presented conflicting results. As has been observed previously and confirmed in this study, clo-lip preclude the development of RA by ablating all circulating monocytes (Misharin et al., 2014; Solomon et al., 2005). The fact that restoring NCM in these mice enables the progression of STIA appears to conflict with sensitivity of NCM-deficient NR4A1^-/-^ mice to arthritis (Brunet et al., 2016) (Liebmann et al., 2018). Our data resolve this conflict by identifying an e.v. Syn Ly6C^-^ population that is unaffected in number or transcriptional profile in NR4A1^-/-^ mice. Through injection of intravenous anti-CD45 antibody, we confirm the extravascular localization of this population. Furthermore, we are able to discriminate between the i.v. and e.v. populations of Syn Ly6C^-^ cells through cell surface expression of FcRγRIV or Treml4 and CD177, respectively. Although e.v. Syn Ly6C^-^ cells are independent of NR4A1, NOD2 may play a role in their development since it has previously been linked to restoring circulating NCM numbers that lack NR4A1 (Lessard et al., 2017). Finally, e.v. Syn. Ly6C^-^ cells are present in synovial biopsies from RA patients and their depletion prevents experimental RA in mice. Taken together, our data suggests that e.v. Syn Ly6C^-^ cells represent a novel population distinct from circulating cells that are necessary for the development of RA.

In order to explore the heterogeneity of tissue Ly6c^-^ myeloid cells but exclude macrophages, we performed single-cell RNA-seq on CD64^-^Tim4^-^ mononuclear phagocytes from the synovium. Analysis of this data enabled us to distinguish a subpopulation of monocyte-derived dendritic cells. Thus, we used MHCII expression in following studies to distinguish between these MHCII^+^ mono-DC and the MHCII^-^ Syn. Ly6c^-^ cells of interest. Based on our results, these cells represent a very small fraction of CD45^+^CD11b^+^Ly6G^-^ murine synovial cells analyzed in single-cell RNA-seq data published by Culemann et al (Culemann et al., 2019). Nevertheless, several of the markers associated with the e.v. Syn. Ly6c^-^ population are reported in the Culemann et al data set, thereby validating our results. Moreover, we observe an overlap between genes differentially expressed in our populations and those described in day 5 post-STIA. These results demonstrate that e.v. Syn. Ly6c^-^ cells can be distinguished from other myeloid cells in the steady-state and arthritic mouse joint.

Further, two recent studies utilized single-cell RNA seq to characterize human myeloid cells from the joints of RA patients (Alivernini et al., 2020; Zhang et al., 2019). However, neither of these studies rigorously discriminated between macrophages and other myeloid cells that were present in the synovium. By using a module score based on marker genes to annotate cells, we identified a small fraction of cells among the limited number of CD14^+^ cells (750) in the AMP study (Zhang et al., 2019) that matched either MHCII^+^ or MHCII^-^ Syn Ly6c-populations from our single-cell data. We observed some differences between OA patients and RA patients classified as either leukocyte-rich or leukocyte-poor but a larger cohort of patients and an expanded single-cell data set is needed to determine whether a true association exists. The Alivernini group performed single-cell RNA-seq on CD64^+^CD11b^+^CD3^-^CD19^-^CD20^-^CD56^-^ CD49^-^CD117^-^CD15^-^ synovial cells from healthy controls as well as from RA patients who are treatment-naïve/resistant or in clinical remission (Alivernini et al., 2020). We found that our MHCII^-^ murine population was most closely associated with MERTK^-^CD48^+^SPP1^+^ synovial myeloid cells, which were significantly enriched active RA patients compared with those in remission or healthy controls (Alivernini et al., 2020). This is consistent with both our proposed role of e.v. Syn Ly6c-cells and the characterization of MERTK^-^CD48^+^SPP1^+^ synovial myeloid cells as activated, glycolytic, pro-inflammatory, and involved in migration. In contrast, our MHCII^+^ murine population was associated with both the MERTK+FOLR+ID2+ and ICAM1+ populations, which were identified as macrophage proliferating precursors and pro-inflammatory cells, respectively. Thus, our novel e.v. Syn Ly6c-cells are likely to be relevant in the context of human disease.

e.v. Syn Ly6C^-^ cells may represent a terminal monocyte population. In addition to lacking canonical macrophage surface markers CD64, MERTK, and TIM4, e.v. Syn Ly6C^-^ express low levels of MafB compared to CD64^+^ synovial macrophages. MafB has been previously associated with tissue macrophages over monocytes (Lavin et al., 2014) and is central for suppressing macrophage proliferation (Kelly et al., 2000). Our experiments determined that e.v. Syn Ly6C^-^ cells maintain their ability to proliferate and do not differentiate into macrophages. The e.v. Syn Ly6C^-^ cells are also negative for the DC master regulator Zbt46 (Meredith et al., 2012; Satpathy et al., 2012) as compared to the mono-DCs. Collectively, these data support e.v. Syn Ly6C^-^ cells as a self-renewing terminal monocyte population that is distinct from DCs and macrophages in the tissue.

Our lineage-tracing studies with CX3CR1^CreER^zsGFP suggest that e.v. Syn Ly6C^-^ are embryonically derived similar to tissue macrophages are GFP^+^, similar to previous studies (Ginhoux and Guilliams, 2016; Gomez Perdiguero et al., 2015a; Gomez Perdiguero et al., 2015b; Guilliams et al., 2013; Mildner et al., 2017; Yona et al., 2013). The incomplete penetrance of GFP in this population may be attributed to the precise timing of tamoxifen injection (E15) which failed to label all progenitors or the loss of GFP as e.v. Syn Ly6C^-^ cells proliferate. Nonetheless, our CX3CR1^CreER^zsGFP bone marrow chimeric mice confirm that e.v. Syn Ly6C^-^ cells are not derived from circulating monocytes. These results are in contrast to i.v. Syn Ly6C^-^ cells which are derived from circulating cells and were GFP^+^ in CX3CR1^CreER^zsGFP bone marrow chimeric mice but not in the lineage-tracing studies. Expansion of e.v. Syn Ly6C^-^ cells is in CCR2^-/-^ and NR4A1^-/-^ mice during STIA, also supports their ability to locally proliferate.

Increased vascularity and enhanced permeability of the synovium has been associated with RA and experimental models of arthritis (Atehortua et al., 2017; Binstadt et al., 2006; Wipke et al., 2002). In steady-state, access to the synovium is restricted by the size of the particle (Cloutier et al., 2012), consistent with failure of the clo-lip to eliminate synovial macrophages. However, since e.v. Syn Ly6C^-^ cells in the synovium are susceptible to depletion by Clo-lip, we proposed that these cells have access to the vasculature through reverse transmigration from synovium to the intra-vascular space. One group demonstrated bidirectional transmigration of monocytes across hepatic sinusoidal endothelium (Zimmermann et al., 2016), and another showed that reverse transmigration contributes to the development of pathogenic foam cells in atherosclerosis (Angelovich et al., 2017). We show that e.v. Syn Ly6C^-^ cells rapidly undergo LFA-dependent reverse transmigration in response to K/BxN serum. LFA1 is required for monocyte crawling (Auffray et al., 2007) and has been implicated in the pathogenesis of arthritis (Suchard et al., 2010; Watts et al., 2005). LFA1^-/-^ mice retain comparable levels of e.v. Syn Ly6C^-^ cells to WT in steady state but this population fails to expand in response to STIA. Moreover, LFA^-/-^ e.v. Syn Ly6C^-^ cells exhibit reduced reverse transmigration and downregulation of genes associated with migration and inflammation compared to controls. These data suggest that LFA is required for e.v. Syn Ly6C^-^ cells proliferation and reverse transmigration in response to arthritic stimuli.

To the best of our knowledge, we are the first to identify a population of tissue-resident Ly6C^-^ cells with distinct functions from DCs or macrophages. e.v. Syn Ly6C^-^ cells respond rapidly to inflammatory signals, drastically expand in numbers, traverse perivascular space via an LFA-dependent mechanism to recruit immune infiltrate for arthritis. Our data support a role for e.v. Syn Ly6C^-^ cells as instigators of synovial inflammation leading to the pathogenic cascade of inflammatory arthritis.

## Acknowledgments

We would like to thank Dr. Steffen Jung for providing critical review of the manuscript. We also like to thank the Northwestern University Lurie Cancer Center Flow Cytometry Core Facility, which is supported by NCI Cancer Center Support Grant P30 CA060553 awarded to the Robert G. Lurie Comprehensive Cancer Center. This research was supported in part through the computational resources and staff contributions provided by the Genomics Compute Cluster, which is jointly supported by the Feinberg School of Medicine, the Office of the Provost, the Office for Research and Northwestern Information Technology. The Genomics Compute Cluster is part of Quest, Northwestern University’s high-performance computing facility, with the purpose to advance research in genomics. SYC was supported by a pre-doctoral AHA award (19PRE34380200). CMC was supported by the Lupus Research Alliance (Novel Research Grant), the Rheumatology Research Foundation (Innovative Research Grant) and the Northwestern University Dixon Translational Research Initiative. HMM was supported by KL2 TR001424, HL134375S1, and AR007611. AVM was supported by NIH grants U19AI135964, P01AG049665, R56HL135124, R01HL153312 and NUCATS COVID-19 Rapid Response Grant. A.B. was was supported by NIH grants HL145478, HL147290, and HL147575. G.R.S.B. was supported by NIH grants U19AI135964, P01AG049665, P01AG04966506S1, R01HL147575 and Veterans Affairs grant I01CX001777. DRW was supported funding from Arthritis National Research Foundation, American Federation for Aging Research, and American Heart Association (18CDA34110224). HP was supported by AR074902, AR075423, CA060553, HL134375, RRF Innovative Research Grant, United States-Israel Binational Science Foundation Investigator Grant, the Precision Medicine Fund, and the Mabel Green Myers Professorship.

## Materials and Methods

### Mice

Breeder pairs were purchased and experimental mice bred in house, and/or acclimated in barrier and specific pathogen-free animal facility at the Center for Comparative Medicine, Northwestern University. Female mice were used for all RA-like studies. All experimental procedures were carried out on mice aged 8-10 weeks (unless stated otherwise in aging studies). To induce serum transfer arthritis, 85_μ_L/20g/mouse was given intravenously (I.V). All procedures were approved by the Institutional Animal Care and Use committee at Northwestern University.

### Flow cytometric analysis

To prepare single-cell suspensions from joint, joints were removed from hind paws following euthanasia, perfused and stored on ice in sterile HBSS. Skin and toes were removed from each paw and bone marrow flushed from exposed tibia with sterile HBSS through a 30G needle. Synovial tissue was then infused with 1.5mL/joint of ankle digestion buffer (2.4mg/mL dispase II, 2mg/mL collagenase D, 0.2mg/mL DNAse I in HBSS pH 7.2-7.6) before incubation at 37°C for 1h with shaking. Cells were then aggravated through a 40-_μ_m mesh filter. Red blood cells were removed with lysis 250_μ_L/sample (1x PharmLyse in sterile water) at room temperature for 1 minute. Dead cells were stained with eFluor 506 viability dye (1:1000 dilution). Cells were incubated with FcBlock (BD Bioscience) and stained with selected antibodies outlined below. Cells were fixed with 10% PFA at 4°C for 20 minutes. To prepare single cell suspensions from blood, 90_μ_L blood collected by cardiac puncture was incubated with FcBlock and selected antibodies. Red blood cells were lysed with FACS lyse at rt for 10 minutes (1x in sterile water) and single-cell suspensions were acquired on BD LSR II or BD Symphony. For all FACSorting studies, cells were acquired on a BD FACSAria. Count eBeads were used in joint preparations to calculate cell numbers. Fluorescence minus one samples were used to set gates. Compensation and analysis of flow-cytometry data was carried out in FlowJo V10.

### Bulk RNA sequencing

RNA from FACSorted synovial cells was extracted using PicoPure RNA Isolation kit as per manufacturer’s instructions. Bulk RNA-seq shown in Figure 1 was carried out using QuantSeq 3’ mRNA sequencing kit, while bulk RNA-seq shown in Figure 4 utilized full-length SMART-seq v4 Ultra Low Input Kit for Sequencing.

Following the sequencing, libraries in the form of BCL files were obtained from Illumina’s BaseSpace platform and demultiplexed (using bcl2fastq v2.17.1.14) to convert them into FastQ read format for further processing. The QuantSeq reads were then processed further by trimming the adapters, low quality bases and short reads (using BBDuk version 37.22 with the following parameters: *k=13 ktrim=r useshortkmers=t mink=5 qtrim=r trimq=10 minlength=20)*. After trimming, remaining reads were aligned to the mouse genome reference mm10 (Mus Musculus / UCSC assembly GRCm38) using STAR (Dobin et al., 2013). Aligned reads in BAM format were mapped to the reference transcriptome (Mus Musculus GRCm38.87) to obtain exon counts and generate gene expression tables using the tool HTSeq (Anders et al., 2015). The SMART-seq reads were trimmed using Trimmomatic (version 0.36), (Bolger et al., 2014) to remove adapter sequences, low quality bases and short reads (minimum length = 20bp). After trimming, remaining reads were aligned to the mm10 genome reference (Mus Musculus / UCSC assembly GRCm38) with Tophat aligner (tophat 2.1.0). The aligned reads in BAM format were mapped to gene exons by HTSeq as above (Anders et al., 2015) using the reference transcriptome GTF file (Mus Musculus GRCm38.87).

All gene expression counts were scaled to read depth using counts per million reads mapped (cpm). To filter out lowly expressed genes, genes with no group mean above 7 cpm in the relevant cell types were excluded from the analysis. Differentially expressed genes (DEG) across multiple cell types were defined as genes with a difference of 2-fold between any two groups. K-means clustering of DEG was carried out in Gene-E. GO enrichment was calculated using Gorilla (Eden et al., 2007; Eden et al., 2009) on each cluster with all DEG as background. Expressed genes, K-means clustering, and GO processes for all datasets are provided in supplemental files for each figure. Volcano plots were generated in R (version 3.3.1). Principal Component Analysis (PCA) and Pearson’s correlation were performed on expressed genes and visualized with R (version 3.3.1). In Figure 1, monocyte populations were compared with CD64^+^ macrophages that were isolated from the same mice and sorted into 4 subpopulations based on the cell surface expression of CX3CR1 and MHCII.

### Single-cell RNA sequencing

RNA libraries for single-cell analysis were prepared using 10x Chromium Single Cell 3’ Solution v3. Reads were processed and aligned to mm10 mouse reference genome using mkfastq and count commands of cellranger 3.1.0 pipeline. Subsequent analyses, including quality control, unsupervised clustering, identification of cluster markers, and visualization of gene expression were carried out using Seurat v3.1 package in R. Samples were individually assessed and filtered based on the number of UMI counts and % mitochondrial reads per cell. To account for technical variability, sample-specific thresholds were used as indicated below.

| Genotype | Total Cells Detected | Median Genes/Cell | Median UMIs/Cell | Min UMI/cell | Max UMI/cell | Max % MT reads/Cell | # cells after filtering |
| --- | --- | --- | --- | --- | --- | --- | --- |
| C57BL/6 | 9447 | 2681 | 9323 | 4000 | 35000 | 7 | 7160 |
| CCR2<sup>-/-</sup> | 4867 | 3184 | 12927 | 5500 | 40000 | 7 | 3621 |
| NR4A1<sup>-/-</sup> | 13375 | 818 | 1494 | 4000 | 38000 | 7 | 2867 |
| LFA1<sup>-/-</sup> | 7908 | 2622 | 8880 | 3500 | 31000 | 5 | 5444 |

For the initial analysis on the C57BL/6 sample, selection of variable genes was performed using the default vst method with nfeatures set to 2000. UMAP dimensionality reduction and unsupervised graph-based clustering were performed with top 16 principal components (PC) and resolution parameter of 0.2. SingleR package v1.0.5 was used to annotate cells with Immgen reference cell types. Differentially expressed genes for each subpopulation were defined log(fold-change)>|0.25| and adjusted p-value<0.05 by Wilcoxon test with Benjamini-Hochberg procedure for False Discovery Rate. Module scores of cell type signatures based on manually selected genes (Supp. Table 2B) were computed using AddModuleScore function with default parameters. Pearson correlations were calculated between the averaged expression profiles of single-cell subpopulations and bulk RNA-seq on monocyte populations (Figure 1). Cell cycle scoring was performed using G2/M and S phase gene sets provided in Seurat, converted to orthologous mouse genes using BioMart R package. Cells in subpopulations 0, 1, and 3 were further classified as either MHCII^+^ or MHCII^−^ by a threshold of 2 on normalized expression of H2-Eb1. Co-expression of H2-Eb1 with MHCII^+/-^ compartment genes was visualized using DimPlot function with blend=TRUE parameter. Integration of C57BL/6 cells with other samples was executed using Seurat anchoring method with 30 CCA dimensions and visualized by UMAP using top 13 (B6 with CCR2^-/-^ and NR4A1^-/-^) and 15 (B6 with LFA1^-/-^) PCs. Subpopulation labels were determined by the majority label of C57BL/6 annotated cells in each cluster. Relative contributions of the integrated samples were calculated by down-sampling to 2000 (B6 with CCR2^-/-^ and NR4A1^-/-^) and 3000 (B6 with LFA1^-/-^) cells for each sample after clustering, with significance determined through chi-square test. GO processes were calculated using GOrilla on genes increased or decreased in expression log_2_(fold-change)>|0.25|, adjusted p-value<0.05 by Wilcoxon test with Benjamini-Hochberg) in LFA1^-/-^ MHCII^-^ compartment compared to C57Bl/6 with 14144 expressed genes as background.

For annotation of individual human myeloid cells from Zhang et al (Zhang et al., 2019), we calculated the module score of the top 10 orthologous marker genes (by fold change) for the MHCII^+^ and MHCII^-^ compartments (Supp. Table 2C). The scores were normalized into the range of -1 and 1 to enable direct comparison between gene modules. Cells were annotated as the cell type with the highest score over 0.5 or as “none” if no module scored above 0.5. In addition, similarity of our clusters with published clusters in Culemann et al, Zhang et al, and Alivernini et al (Alivernini et al., 2020; Culemann et al., 2019; Zhang et al., 2019) was determined by calculating the fraction of their top 20 reported markers that overlapped with the list of differentially expressed genes for each of our subpopulations 2, 4, and 5 (Supp. Table 2A) as well as MHCII^+^ and MHCII^-^ relative to all other cells (Supp Table 2C). Marker genes were converted into mouse orthologs in the case of the latter two human data sets (Supp. Table 2D).

Significance of overlap was determined by hypergeometric distribution with the 2000 variable genes as background and a FWER cutoff of 0.05 was applied based on Bonferroni correction using the number of comparisons for each data set. To assign C57BL/6 myeloid cells to either i.v. NCM. mono-DC, or e.v. NC, singleR was run with the bulk RNA-seq on these populations as reference.

### Intra and extra vascular labeling of immune cells

To label intra-vascular immune cells, anti-CD45 BUV661 antibody was administered I.V. at 6_μ_g/mouse in 200_μ_l sterile PBS. Mice were then returned to housing environment for 5 minutes before euthanasia or for 1 hour before administration of second I.V. anti-CD43 BUV395 antibody. In studies using STIA, K/BxN serum was administered 5 minutes after anti-CD45 BUV661 antibody, 60 minutes prior to anti-CD43 BUV395 antibody as previously described.

### Monocyte depletion

For depletion studies, 200_μ_l/mouse clodronate-laden liposomes were given I.V. 24 hours prior to euthanasia. All mice were perfused with 20mL with ice-cold HBSS following euthanasia to remove circulating cells and retain adherent intra-vascular cells.

### Bone marrow chimeras

Bone marrow chimeras were generated using C57Bl/6, CX3CR1^CreER^.zsGFP and/or B6.CD45.1 mice. Recipient mice were maintained on autoclaved water supplemented with antibiotics for 2 weeks prior to and 2 weeks post irradiation. Recipient mice were irradiated with 1000 cGy _γ_-radiation, 24 hours before I.V. transfer of 10 million donor bone marrow cells. For shielded chimeras, hind joints were shielded with lead during irradiation and mice were treated with Ketamine (100 mg/kg) and Xylazine (16 mg/kg) prior to irradiation to prevent movement of joints. 6 hours after shielded irradiation mice received 30mg/kg busulfan to deplete remaining bone marrow in shielded joints, before transfer of bone marrow cells as before. To determine the proliferation of monocytes, chimeric mice were treated with 2mg EdU on experimental D0 and 0.5mg EdU for subsequent 14 days. Fate-mapping studies were carried out as previously described (Yona et al, 2013). Briefly, 50mg/kg tamoxifen and 10mg/mL progesterone were dissolved in corn oil and administered by oral gavage at E15 of gestation to pregnant CX3CR1^CreER^ mice. For Cre induction in adult mice, 50mg/kg tamoxifen in corn oil was administered I.P on D0 and D1. For STIA studies using CX3CR1^CreER^.zsGFP, 110μL/20g of K/BxN serum was administered to tamoxifen-treated and naive mice to compensate for immunosuppressive effects of tamoxifen. Disease was assessed using clinical scoring 3 times per week for 21 days. Mice were euthanized at 1h and 24 h after serum injection.

### Statistical analysis

All statistical analysis was carried out in GraphPad Prism V8. P-values less than 0.05 were considered statistically significant using two-tailed unpaired t-test with equal variance.

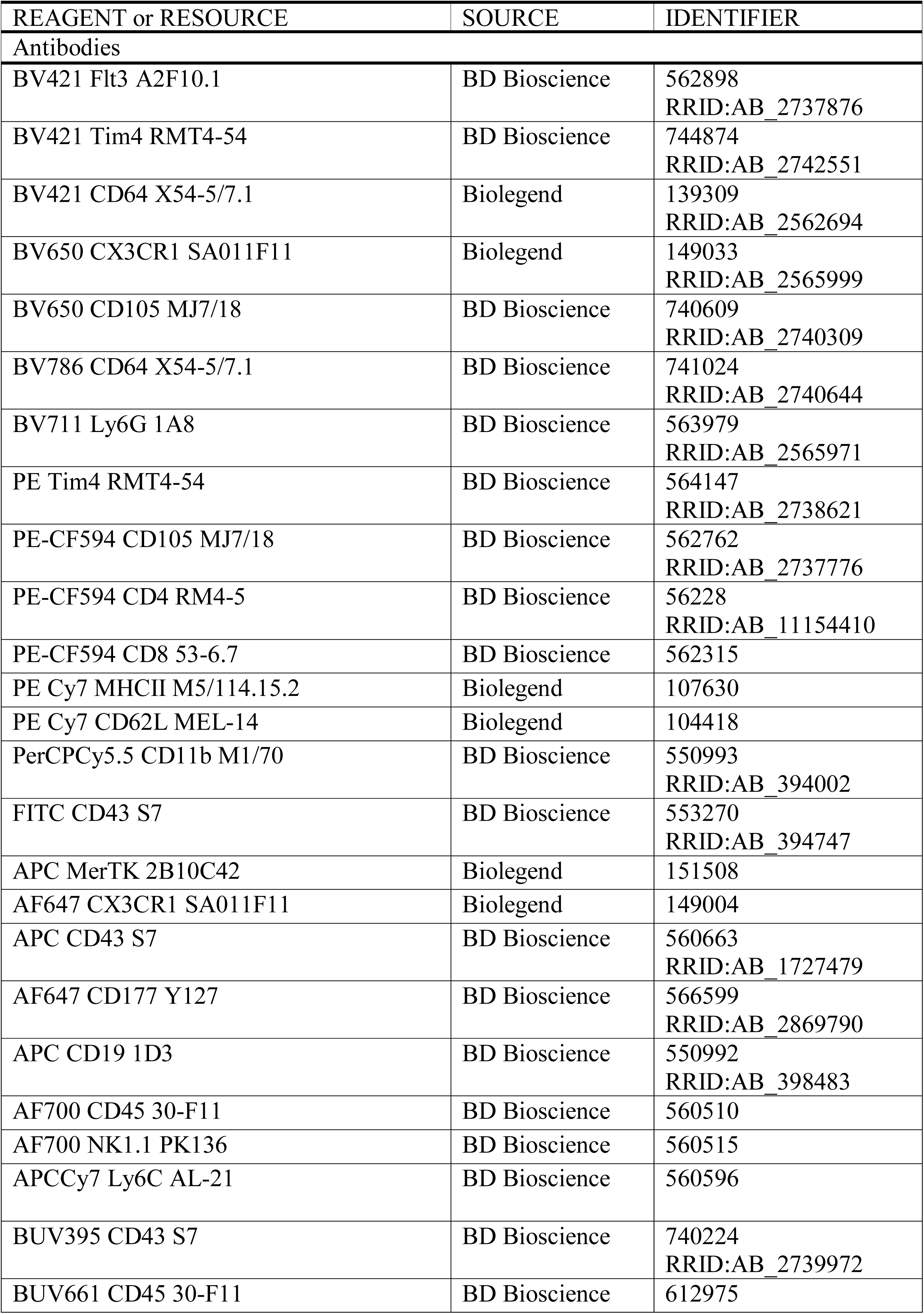

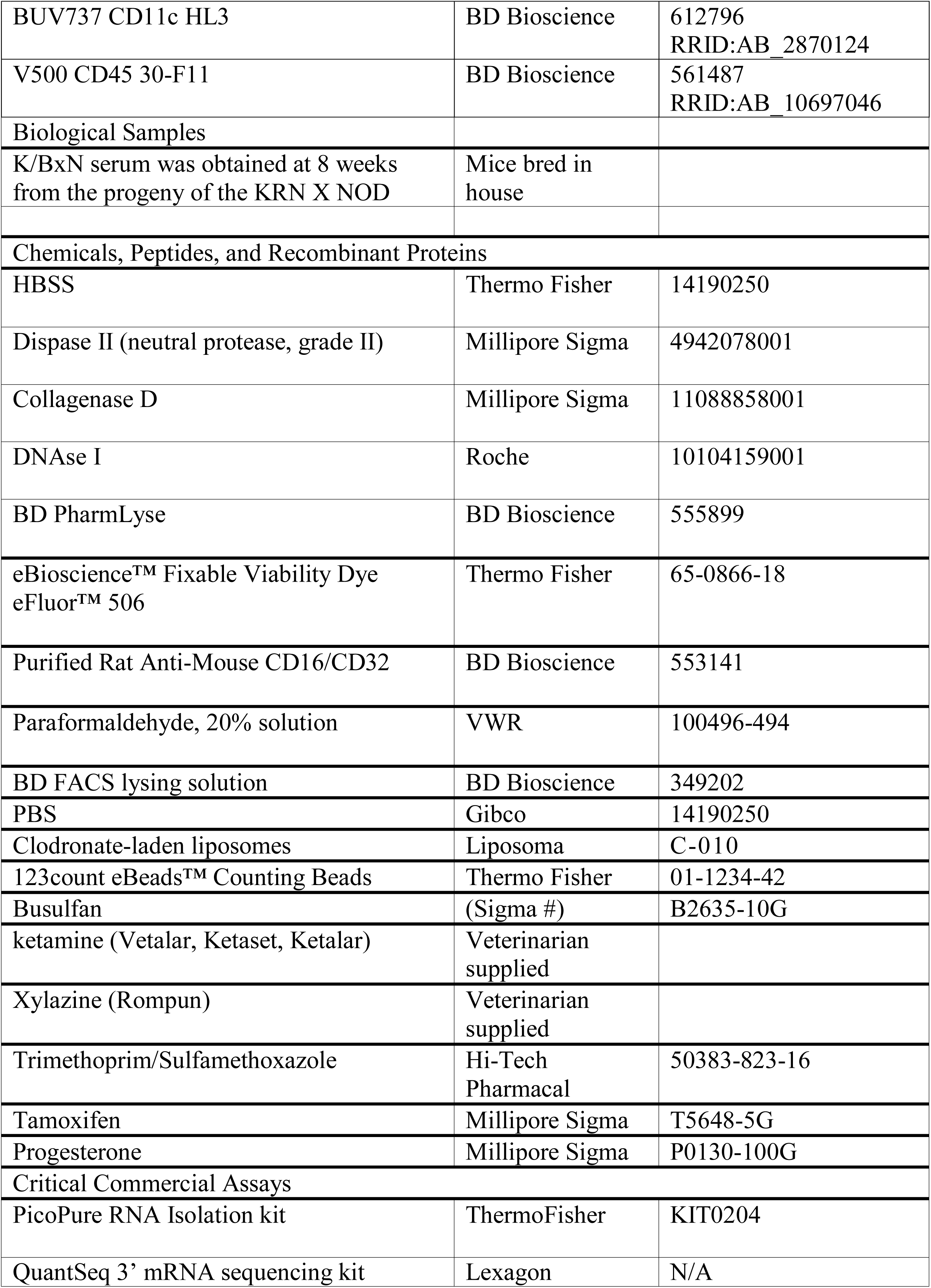

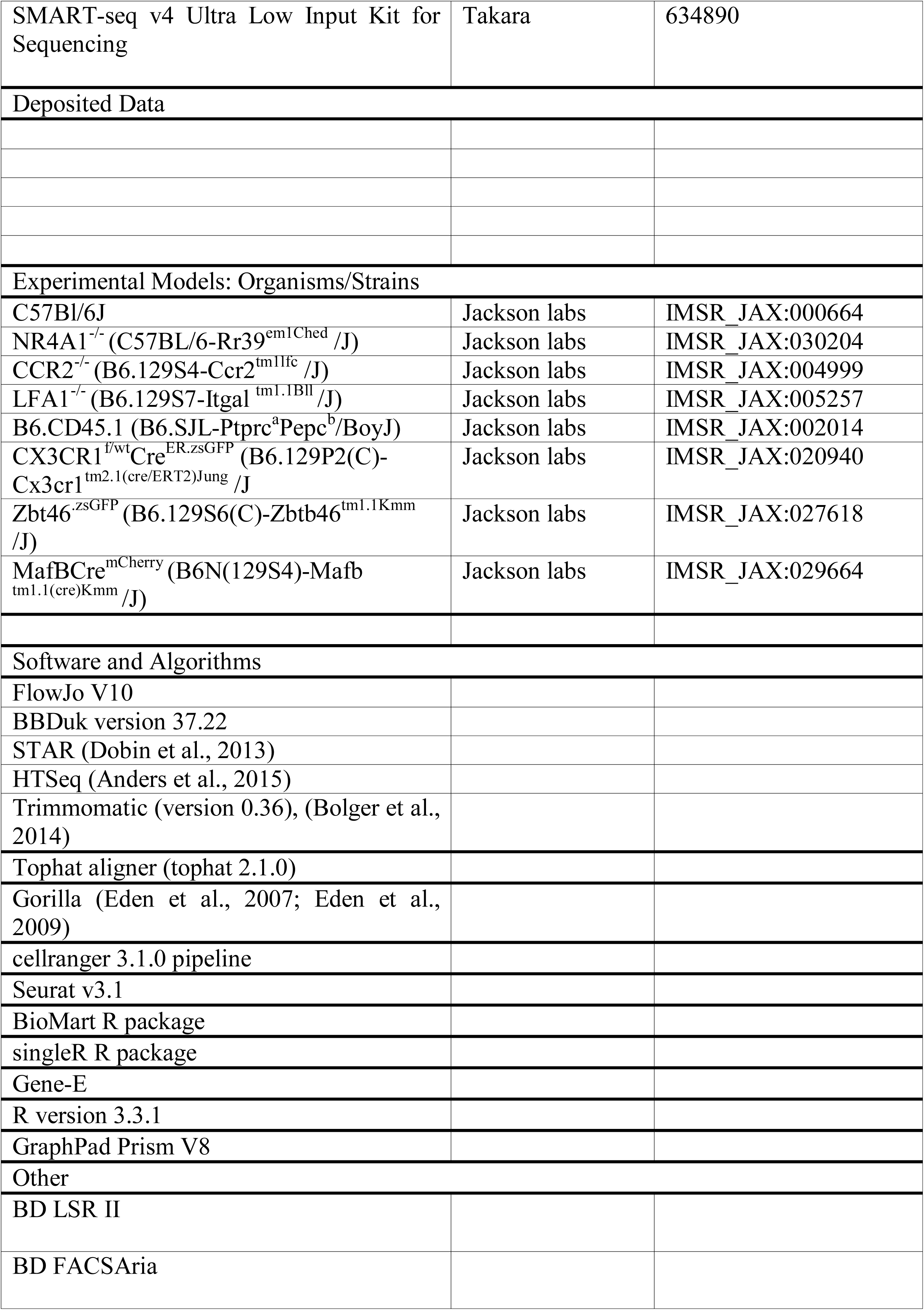

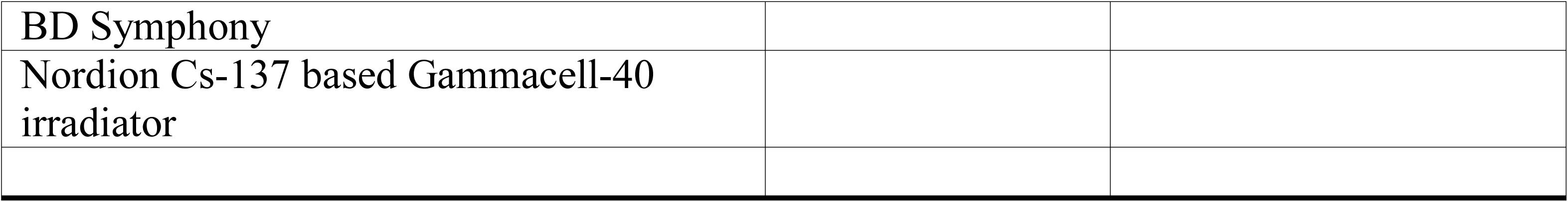

