## Supplemental Figures for "Tissue-resident, extravascular CD64^-^Ly6C^-^ population forms a critical barrier for inflammation in the synovium"

**Supplemental Figures and Tables**

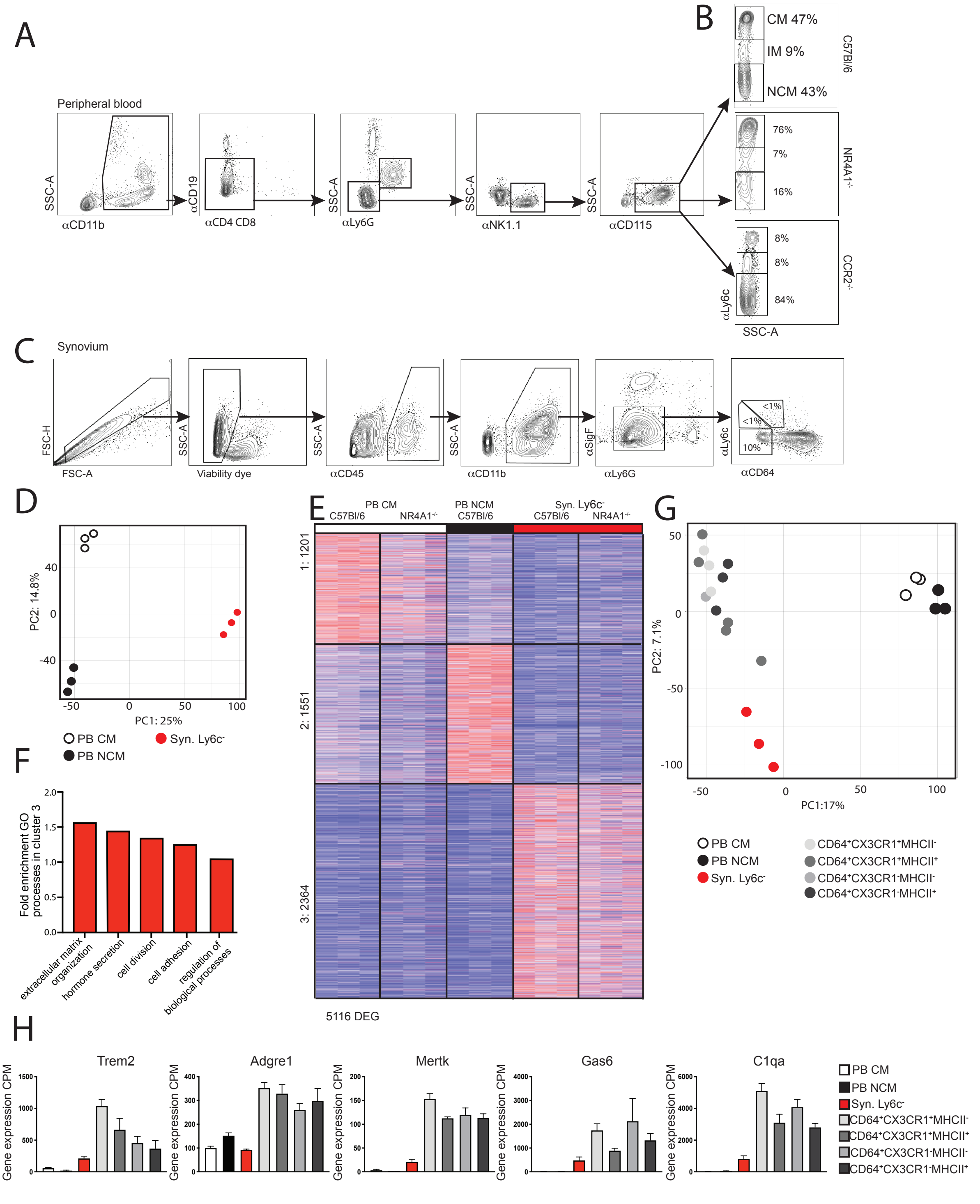


**Supplementary Figure 1:** (A) Flow gating strategy for identification of PB monocyte subpopulations in C57Bl/6, B) NR4A1$^{-/-}$ and CCR2$^{-/-}$ mice, and C) Flow gating strategy for identification of Syn monocytes (CD64$^{-}$) and macrophages (CD64$^{+}$). D) PCA of 10206 genes expressed by PB CM, PB NCM, and Syn Ly6C$^{-}$. E) Visualization of clusters from Figure 1L with expression in PB CM and Syn Ly6C$^{-}$ from NR4A1$^{-/-}$ mice. F) Significantly enriched GO processes in cluster 3 from Figure 1L with preferential expression in Syn Ly6C$^{-}$ ($P<0.05$). G) PCA of 9661 genes expressed in PB CM, PB NCM, Syn Ly6C$^{-}$ and Syn Mac. H) Mean expression of representative macrophage genes from PB CM, PB NCM, Syn Ly6C$^{-}$ and Syn Mac populations (RNA-seq data: n=3, error bars indicate SEM).

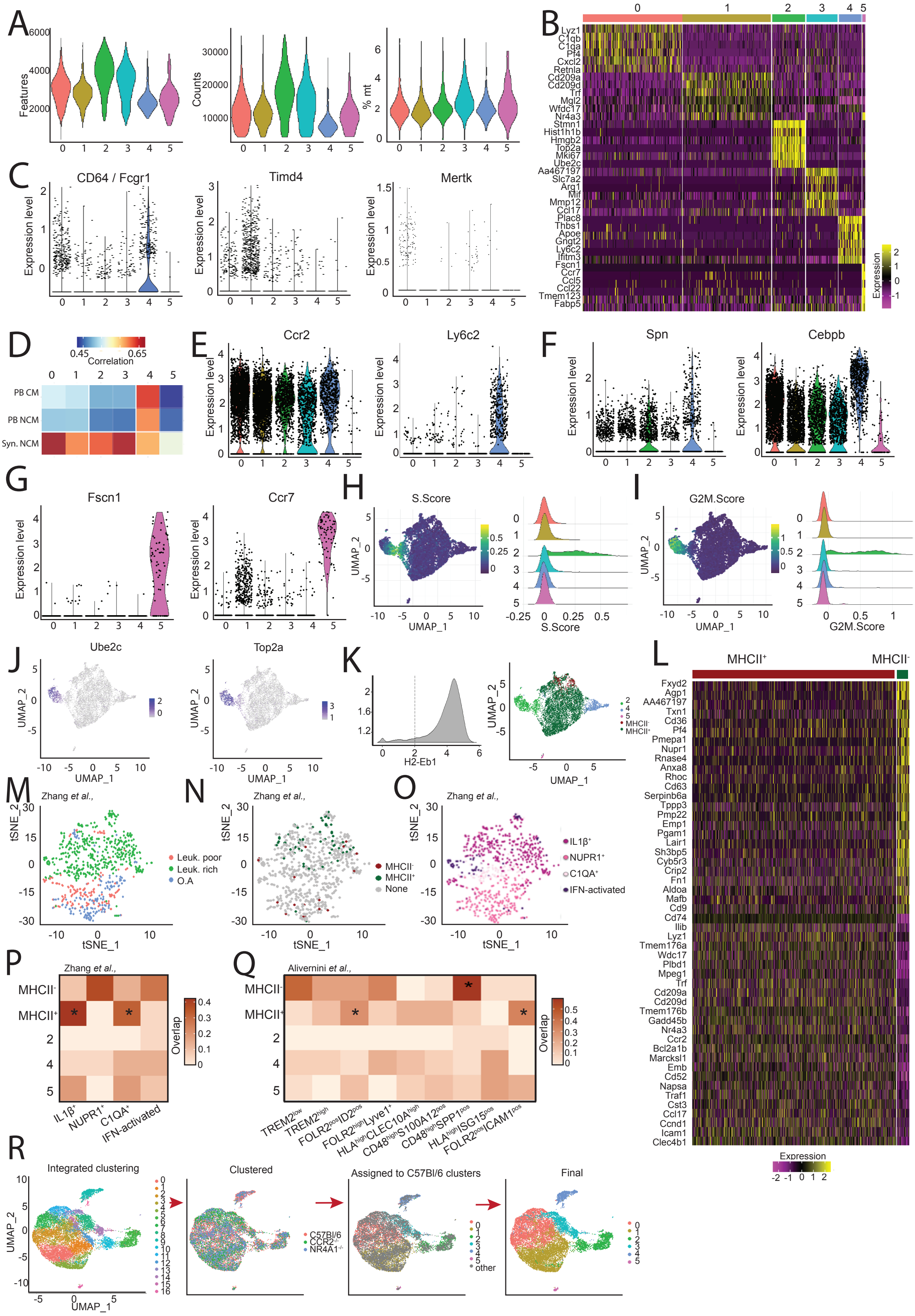
A
Features
Counts
% mt
B
Lyz1
C1qb
C1qa
Pf4
Cxcl2
Retnla
Cd209a
Cd209d
Trf
Mgl2
Wfdc17
Nr4a3
Stmn1
Hist1h1b
Hmgb2
Top2a
Mki67
Ube2c
Aa467197
Slc7a2
Arg1
Mif
Mmp12
Ccl17
Plac8
Thbs1
Apoe
Gngt2
Ly6c2
Ifitm3
Fscn1
Ccr7
Ccl5
Ccl22
Tmem123
Fabp5
Expression
C
CD64 / Fcgr1
Timd4
Mertk
Expression level
D
Correlation
PB CM
PB NCM
Syn. NCM
E
Ccr2
Ly6c2
F
Spn
Cebpb
G
Fscn1
Ccr7
H
S.Score
UMAP_1
UMAP_2
I
G2M.Score
J
Ube2c
Top2a
K
H2-Eb1
MHCII-
MHCII+
L
MHCII+
MHCII-
Fxyd2
Agp1
AA467197
Txn1
Cd36
Pf4
Pmepa1
Nupr1
Rnase4
Anxa8
Rhoc
Cd63
Serpinb6a
Tppp3
Pmp22
Emp1
Pgam1
Lair1
Sh3bp5
Cyb5r3
Crip2
Fn1
Aldoa
Mafb
Cd9
Cd74
Ilib
Lyz1
Tmem176a
Wdc17
Plbd1
Mpeg1
Trf
Cd209a
Cd209d
Tmem176b
Gadd45b
Nr4a3
Ccr2
Bcl2a1b
Marcksl1
Emb
Cd52
Napsa
Traf1
Cst3
Ccl17
Ccnd1
Icam1
Clec4b1
Expression
M
Zhang et al.,
tSNE_1
tSNE_2
Leuk. poor
Leuk. rich
O.A
N
MHCII-
MHCII+
None
O
IL1β+
NUPR1+
C1QA+
IFN-activated
P
Zhang et al.,
MHCII-
MHCII+
Overlap
IL1β+
NUPR1+
C1QA+
IFN-activated
Q
Alivernini et al.,
TREM2low
TREM2high
FOLR2posID2pos
FOLR2highLyve1+
HLAhighCLEC10Ahigh
CD48highS100A12pos
CD48highSPP1pos
HLAhighISG15pos
FOLR2posICAM1pos
R
Integrated clustering
Clustered
C57Bl/6
CCR2-/-
NR4A1-/-
Assigned to C57Bl/6 clusters
other
Final

**Supplemental Figure 2**: A) Quality control of $CD45^{+}CD11b^{+}Tim4^{-}CD64^{-}$ cells by subpopulation from C57Bl/6 mice showing number of genes, number of UMIs, and percent mitochondrial reads per cell. B) Relative expression of top 5 marker genes (by fold change) across subpopulations 0-5. C) Normalized expression of macrophage-associated genes across clusters. D) Pearson's correlation between gene expression in scRNA-seq subpopulations and bulk RNA-seq of PB CM, PB NCM, and Syn $Ly6c^{-}$. E) Normalized expression of CM, F) NCM and G) DC associated genes. H) Normalized expression of S-phase and I) G2-phase module genes. J) Visualization of expression of cell cycle genes. K) Re-classification of clusters 0, 1, and 3 as $MHCII^{+}$ and $MHCII^{-}$ based on expression of H2-eb1. L) Relative expression of top 20 differentially expressed genes between the $MHCII^{+}$ and $MHCII^{-}$ cells. M) Annotation of human synovial myeloid cells in (Zhang et al., 2019) by disease phenotype of patient of origin, N) assignment to $MHCII^{+}$ and $MHCII^{-}$ subpopulation based on module score of top 10 marker genes, or O) cluster defined in published AMP data. P) Fraction overlap of differentially expressed genes from C57Bl/6 scRNA-seq subpopulations with top 20 markers of human synovial myeloid cells in (Zhang et al., 2019) and Q) (Alivernini et al., 2020). * indicates significant p-value by hypergeometic test after FWER correction R) Integration of scRNA-seq data from $CCR2^{-/-}$ and $NR4A1^{-/-}$ mice with superimposed C57Bl/6 annotations.

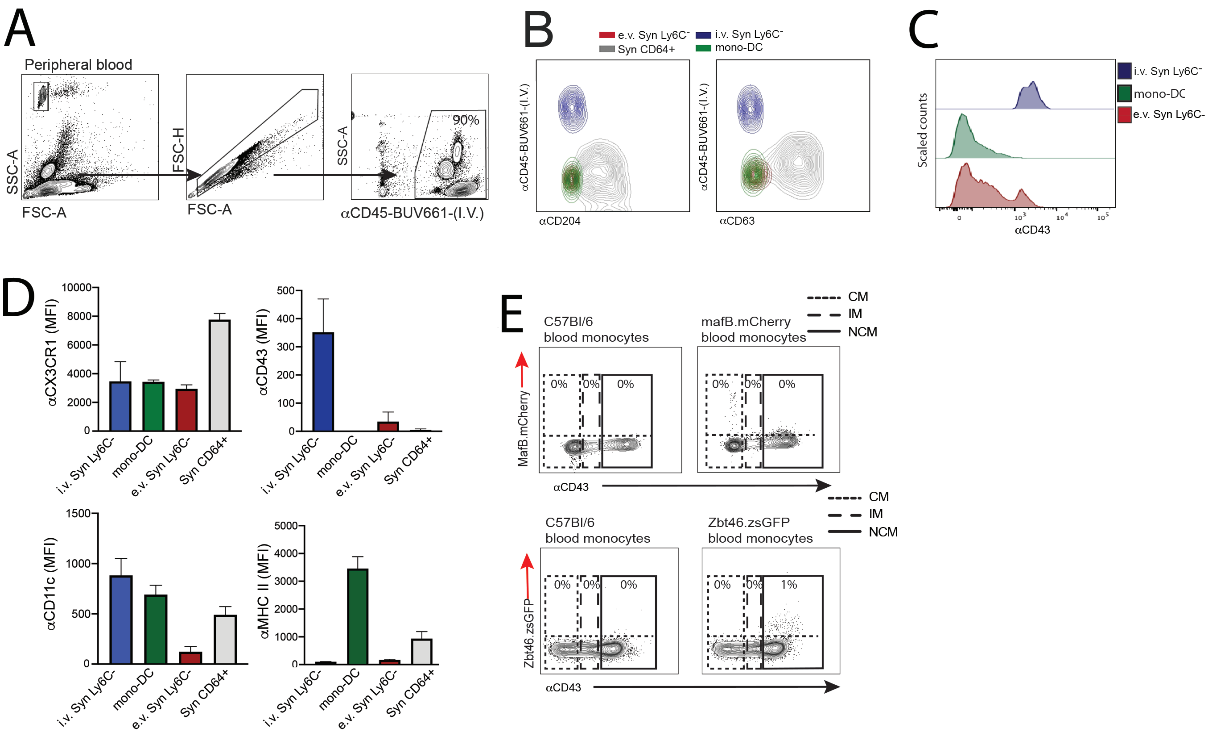


**Supplemental Figure 3**: A) I.V. labeling of cells in peripheral blood using i.v. anti-CD45 antibody. B) Expression of CD204 and CD63 and C) CD43. D) median fluorescent intensity (MFI) of CX3CR1, CD43, CD11c and MHCII in i.v. Syn Ly6C$^-$, mono-DC, e.v. Syn Ly6C$^-$ and Syn CD64$^+$ cells E) Expression of MafB and ZBT46 in PB monocytes of fluorescent reporter mice. Bar graphs are mean N>4+SEM.

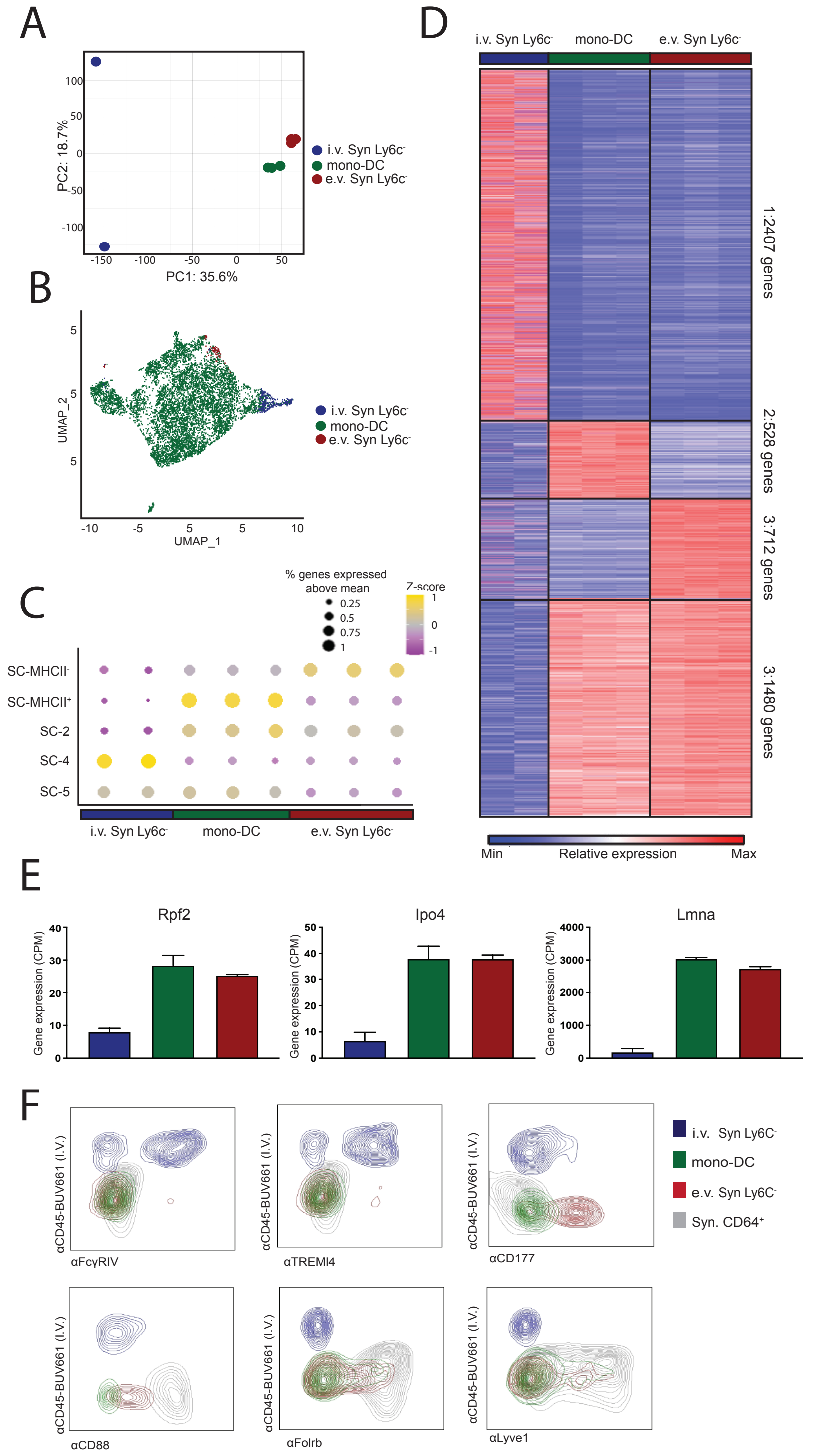


**Supplemental Figure 4:** A) PCA of 10270 expressed genes from i.v. Syn Ly6C$^-$, mono-DC and e.v. Syn Ly6C$^-$ cells. B) Assignment of scRNA-seq cells (Figure 2) to i.v. Syn Ly6C$^-$, mono-DC or e.v. Syn Ly6C$^-$ cells using the bulk transcriptional data as reference in singleR. C) Expression of marker gene sets from single-cell RNA-seq subpopulations in bulk populations. Color of circle indicates z-score normalized expression while size indicates % of genes in the set expressed in the given sample above their mean value. D) K-means clustering of 5127 differentially expressed genes across i.v. Syn Ly6C$^-$ cells, mono-DC and e.v. Syn Ly6C$^-$ cells. E) Example genes with preferential expression in both e.v. Syn Ly6C$^-$ cells and mono-DC. F) Contour plots of αCD45-BUV661 (I.V. CD45 vs αFcγRIV, αTREMl4, αCD77, αCD88, αFolrb, and αLyve1). (RNA-seq data: n=2-3, error bars indicate SEM).

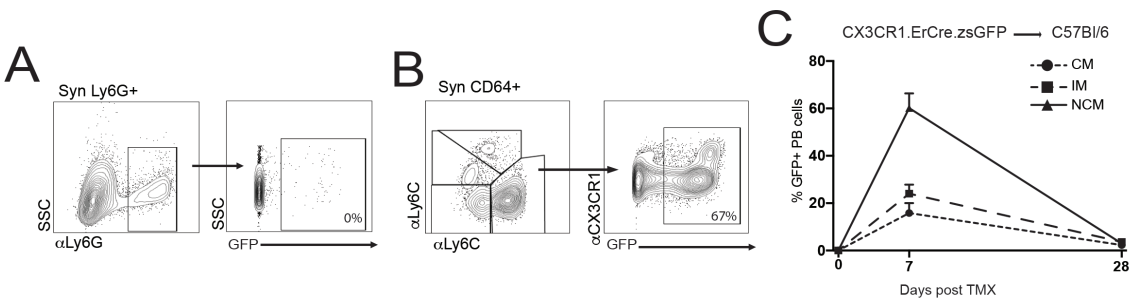


**Supplemental Figure 5:** A) Expression of GFP in synovial Ly6G$^{+}$ and B) CD64$^{+}$ in adult CX3CR1$^{Cre.ER}$.zsGFP mice treated with 50mg/kg TMX at D-1, and D0 analyzed on D1. C) GFP$^{+}$ cells in PB monocytes in CX3CR1$^{Cre.ER}$.zsGFP -> C57Bl/6 bone marrow chimeric mice 7 and 28 days post TMX. All graphs are mean N>4.

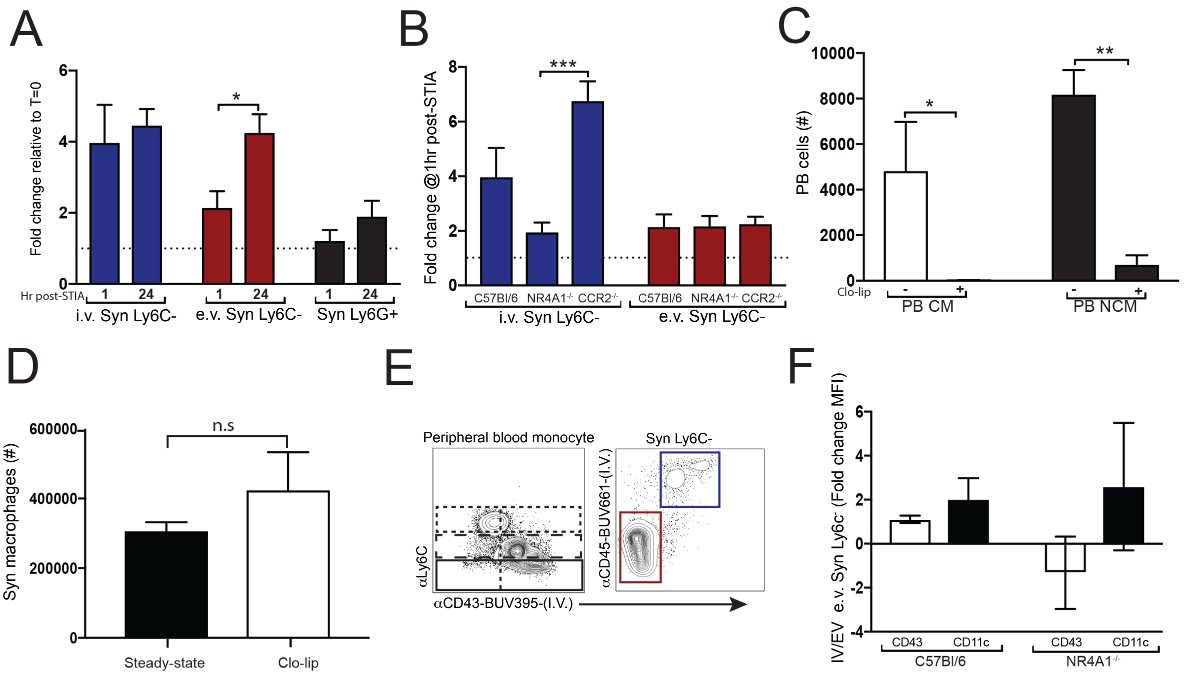


**Supplemental Figure 6**: A) Fold change of i.v. Syn Ly6C$^{-}$ cells, e.v. Syn Ly6C$^{-}$ cells and synovial Ly6G$^{+}$ at 1h and 24h after STIA compared to steady state. B) Fold-change of i.v. Syn Ly6C$^{-}$ cells and e.v. Syn Ly6C$^{-}$ cells numbers 1h after STIA in C57Bl/6, NR4A1$^{-/-}$, and CCR2$^{-/-}$ mice. C) Depletion of PB monocytes and D) synovial CD64+ macrophages following treatment with 200µL of Clo-lip. E) Expression of Ly6C vs αCD43-BUV395 staining in PB and synovial Syn Ly6C$^{-}$ cells. F) Fold-change of CD43 and CD11c of I.V. αCD43-BUV395$^{+/-}$ e.v. NCM in C57Bl/6 and NR4A1$^{-/-}$ cells. All graphs are mean N>4 +SEM P-value was calculated with unpaired t-test. *=p<0.05, **=p<0.01, ***=p<0.005.

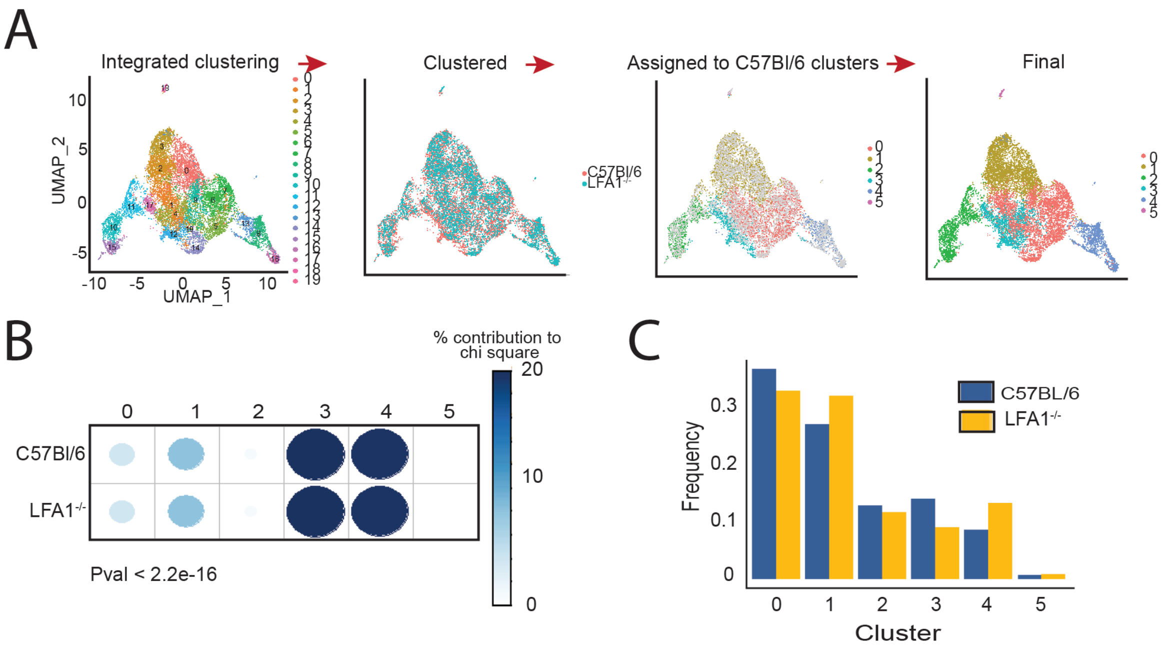


**Supplemental Figure 7**: A) Integration of scRNA-seq data from LFA1$^{-/-}$ mice with superimposed C57Bl/6 annotations. B-C) Proportion of cells annotated as each subpopulation in LFA1$^{-/-}$ and C57Bl/6 mice and chi-squared residuals.
